# A haplotype-resolved reference genome of *Quercus alba* sheds light on the evolutionary history of oaks

**DOI:** 10.1101/2024.02.13.579671

**Authors:** Drew A. Larson, Margaret E. Staton, Beant Kapoor, Nurul Islam-Faridi, Tetyana Zhebentyayeva, Shenghua Fan, Jozsef Stork, Austin Thomas, Alaa S. Ahmed, Elizabeth C. Stanton, Allan Houston, Scott E. Schlarbaum, Matthew W. Hahn, John E. Carlson, Albert G. Abbott, Seth DeBolt, C. Dana Nelson

## Abstract

- White oak (*Quercus alba*) is an abundant forest tree species across eastern North America that is ecologically, culturally, and economically important.
- We report the first haplotype-resolved chromosome-scale genome assembly of *Q. alba* and conduct comparative analyses of genome structure and gene content against other published Fagaceae genomes. In addition, we probe the genetic diversity of this widespread species and investigate its phylogenetic relationships with other oaks using whole-genome data.
- Our genome assembly comprises two haplotypes each consisting of 12 chromosomes. We found that the species has high genetic diversity, much of which predates the divergence of *Q. alba* from other oak species and likely impacts divergence time estimation in *Quercus*. Our phylogenetic results highlight phylogenetic discordance across the genus and suggest different relationships among North American oaks than have been reported previously. Despite a high preservation of chromosome synteny and genome size across the *Quercus* phylogeny, certain gene families have undergone rapid changes in size including resistance genes (R genes).
- The white oak genome represents a major new resource for studying genome diversity and evolution in *Quercus* and forest trees more generally. Future research will continue to reveal the full scope of genomic diversity across the white oak clade.

## Introduction

Oaks (*Quercus* spp.) are important members of ecosystems throughout much of the world (Kremer & Hipp, 2020). In eastern North America, white oak (*Quercus alba*) is often considered a keystone species and is one of the most abundant forest trees across much of its range (Rogers, 1990; Fralish, 2004). In addition to its ecological and cultural importance (Abrams, 2003; Bocsi *et al*., 2021b,a; Stringer & Morris, 2022), white oak has significant economic importance, including a number of high-value timber applications and as the primary species used to cooper barrels for aging distilled spirits (Stringer & Morris, 2022; Dhungel *et al*., 2023). However, few studies have addressed the genomic diversity of *Q. alba*, and a lack of available genetic and genomic resources currently present barriers to furthering understanding of white oak biology and evolutionary history.

*Quercus* (Fagaceae) comprises ca. 500 species, often divided into two subgenera: *Cerris* and *Quercus* (Hipp *et al*., 2020). The latter is typically further divided into the white oaks (section *Quercus*), to which *Q. alba* belongs, and the red oaks (section *Lobatae*). The phylogeny of oaks has been the focus of several recent studies utilizing reduced representation genome sequencing (Sork *et al*., 2016; Hipp *et al*., 2020; Manos & Hipp, 2021), which have clarified some relationships within section *Quercus*.

However, phylogenetic inference in oaks is likely complicated by the suggested prevalence of hybridization and introgression in the group (McVay *et al*., 2017; Lazic *et al*., 2021).

The first published oak genome was that of *Quercus robur* (Plomion *et al*., 2016a), the pedunculate oak, which is common throughout western Eurasia. To date, there have been at least nine *Quercus* species with published chromosome-scale genomes, including four annotated genomes from the white oak clade (Plomion *et al*., 2016a; Han *et al*., 2022; Liu *et al*., 2022; Sork *et al*., 2022; Zhou *et al*., 2022; Ai *et al*., 2022; Kapoor *et al*., 2023; Wang *et al*., 2023). This growing number of annotated genomes allows for comparative analyses of gene content and inferences of genome evolution across the oak phylogeny.

Disease resistance-related genes (R genes) have been a focus of genomic studies on oaks and other tree species because of their central role in plant immunity to pathogens (Plomion *et al*., 2018; Sork *et al*., 2022; Ai *et al*., 2022). Resistance genes confer defense against various viral, bacterial, and eukaryotic pathogens by encoding proteins that recognize pathogen-related molecules and trigger downstream immune responses. Plomion *et al*. (2018) suggested that an expansion of R genes might be at least partly responsible for allowing tree species to live for multiple centuries. Ai *et al*. (2022) found that the *Q. mongolica* genome assembly contained far fewer putative R genes compared to the earlier genome assemblies of *Q. lobata*, *Q. robur*, and *Q. suber*. However, the history of how these R gene families have evolved across the oak phylogeny remains poorly understood.

In addition to providing insights into fundamental questions about plant evolution, improving understanding of genomes across the white oak clade may benefit tree breeding and genetic improvement efforts and help land managers plan for and address global change. Although white oak is generally thought to be more resistant to emerging diseases such as sudden oak death and oak wilt than species in the red oak clade, *Q. alba* does face declining seedling recruitment in many parts of its range (Dhungel *et al*., 2023), which may have implications for ecosystem function throughout eastern North America.

Furthermore, anthropogenic climate change is causing a mismatch between populations and their historical climates (Piao *et al*., 2019; Kijowska-Oberc *et al*., 2020). The amount of standing genetic variation and the extent to which populations are locally adapted will have implications for the response of *Q. alba* and other white oak species to global climate change.

Here, we report the first assembled genome of *Q. alba* and use this new resource to study the evolution of oak genomes. Specifically, we address the following questions: 1) What is the extent of genetic diversity and population differentiation within *Q. alba*? 2) Are previous phylogenetic hypotheses for the relationships among oak species supported by whole genome data? and 3) How have gene content and disease resistance genes evolved during the history of *Quercus* and related taxa? The answers to these questions will provide a roadmap for future work on the white oak group as a model clade for studying genome evolution and adaptation in highly outcrossing forest trees.

## Materials and Methods

### Reference Genome

An individual of *Q. alba* in a forest stand near Loretto, Kentucky, USA (37° 39.0583’, -85° 21.2615’), referred to as MM1 (**Figure 1**), was sampled with the permission of the landowners. Following high-molecular weight DNA extraction and sequencing, PacBio Sequel II HiFi and Hi-C (Phase Genomics) data were assembled with Hifiasm v.0.16.1 (Cheng *et al*., 2021) and scaffolded with 3D-DNA v201008 (Dudchenko *et al*., 2017**; Methods S1**). Two resolved haplotypes were produced, herein referred to as hapA and hapB. The plastome and mitochondrial genomes of MM1 were assembled and annotated separately (**Methods S1**). To assess genome quality, BUSCO v5.2.2 and the embryophyta_odb10 database were used to analyze the hapA and hapB assemblies (Manni *et al*., 2021). The synteny and structural variation of the haplotypes were analyzed with the Synteny and Rearrangement Identifier (SyRI) v1.5.4 (Goel *et al*., 2019) based on a mapping of hapB to hapA with minimap2 v2.24 (Li, 2018). Visualizations of the SyRI results were constructed with plotsr v0.5.1 (Goel & Schneeberger, 2022).

**Figure 1.**
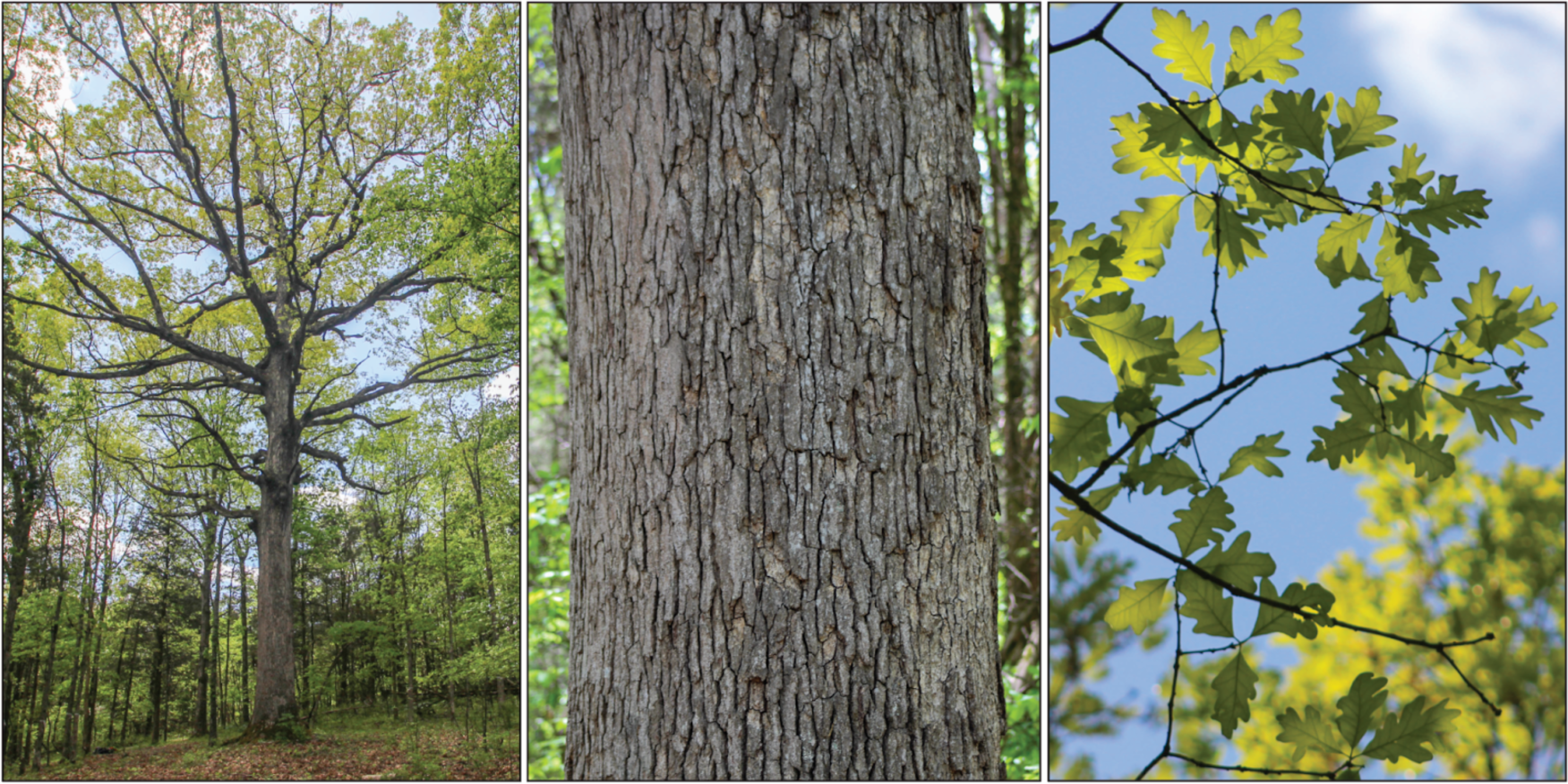
MM1, the *Q. alba* individual sequenced for the genome assembly, growing at Star Hill Farm, Loretto, KY, USA. Photo attribution: D. Larson.

Repetitive DNA and gene annotation were completed independently for the hapA and hapB assemblies (**Methods S1**). BRAKER2 and TSEBRA were used to annotate the hapA and hapB assemblies (Bruna *et al*., 2021; Gabriel *et al*., 2021). RNA sequencing was performed on several tissue types from four individuals and analyzed alongside RNA data obtained from NCBI (SRR006309 – SRR006312) (**Methods S1**).

A genetic linkage map was constructed based on 184 full- and half-sib *Q. alba* individuals (**Methods S1**). Genetic markers with available sequences were aligned to the chromosomes and unplaced scaffold sequences with BLAST+ (Camacho *et al*., 2009). Visualization of the alignment was produced by RIdeogram (Hao *et al*., 2020), excluding markers with multiple equivalent e-value hits at different locations.

The locations of rDNA arrays in hapA and hapB were identified with rnammer (Lagesen *et al*., 2007). Fluorescence *in situ* hybridization (FISH) with rDNA oligonucleotide probes was conducted following the methods of Kapoor et al. (2023). R gene domains and categories were determined using DRAGO2 (Calle García *et al*., 2022). Mechanisms of gene duplication were inferred by the MCScanX duplicate gene classifier tool (Wang *et al*., 2012) of the hapA assembly against itself, resulting in the assignment of five categories of gene expansion type: (1) whole genome/segmental duplication where a set of collinear genes are found duplicated in collinear blocks, (2) tandem duplication where duplicate genes are adjacent to each other, (3) proximal duplication where genes are not adjacent but are in nearby chromosomal regions, (4) dispersed duplications where duplicated genes are found but do not fit the criteria for other categories, and (5) single copy (‘singleton’) genes that have no identified duplication from other genes.

### Population genomics

We sampled 16 *Q. alba* individuals (four individuals each of Wisconsin, Ohio, Indiana, and Mississippi provenances), growing as part of a provenance trial near Vallonia, Indiana, as well as two additional *Q. alba* individuals from Kentucky (**Table S1; Methods S1**). We also included the MM1 tree in our population genetic analyses – the sequence data utilized for this individual were the two runs of PacBio HiFi long reads described above. Sequencing data were analyzed with GATK (McKenna *et al*., 2010; DePristo *et al*., 2011; Van Der Auwera *et al*., 2013) following read trimming and processing (Li & Durbin, 2009; Li *et al*., 2009; Andrews, 2010; Jiang *et al*., 2014; Sim *et al*., 2022; Broad Institute, 2023) to produce a “filtered SNP dataset” used in several downstream analyses. To conduct genetic clustering with *Structure* v2.3.4 (Pritchard *et al*., 2000), 10,000 SNPs were randomly selected from the filtered SNP dataset after additional processing (**Methods S1**). We summarized our *Structure* results with the CLUMPAK online server and default settings (Kopelman *et al*., 2015). We also conducted a principal component analysis (PCA) with PLINK v2.0 (Purcell *et al*., 2007) and visualized the first two principal components, which correspond to the first two eigenvectors, using the ggplot2 library in R (R Core Team, 2013; Wickham, 2016). Pairwise *F*ST values among “populations” were calculated by grouping individuals in three ways **(Methods S1)**, using both the Weir & Cockerham (1984) and Hudson *et al*. (1992) estimators on a per-SNP basis with PLINK v2.0 and then calculating genome wide averages.

Genome-wide nucleotide diversity (π) was calculated using VCFtools v0.1.13 (Danecek *et al*., 2011; Methods S1). <u>Phylogenomics</u> A phylogenomic dataset was assembled using data from several sources including new sequencing of seven species of section *Quercus* and NCBI (**Table S1; Methods S1**). We called variants using the *mpileup*, *call*, and *consensus* commands and the hapA reference in BCFtools (Li, 2011) to produce whole-genome pseudo-reference sequences for each sample, masked repetitive DNA, and randomly selected one allele at heterozygous sites. As a final step to prepare our pseudo-reference sequences, we used the Referee package (Thomas & Hahn, 2019) to calculate genotype quality scores for each site and masked sites where the final base call was not supported (**Methods S1**). To infer phylogenetic relationships, we generated a matrix of whole genome alignments (i.e. all 12 chromosomes, excluding unplaced scaffolds) which included 37 individuals from 19 species of *Quercus* and one individual of *Lithocarpus* as an outgroup. A maximum likelihood phylogenetic tree was estimated with IQ-TREE v2.2.0; site concordance factors (sCF) were calculated with the “--scf” option and 1000 quartet replicates (Minh *et al*., 2020b,a; Mo *et al*., 2023). To investigate the shared genetic variation between *Q. alba* and other oak species we used the same pseudo-reference matrix **(Methods S1).** An ultrametric phylogeny was generated with branch lengths scaled to time using the penalized likelihood approach as implemented in treePL (Sanderson, 2002; Smith & O’Meara, 2012) and a calibration for the crown age of *Quercus* at 56 Ma (Hofmann, 2010; Hofmann *et al*., 2011; Hipp *et al*., 2020), after accounting for ancestral polymorphism by using our estimate of π in *Q. alba* (Edwards & Beerli, 2000; **Methods S1)**.

We also generated phylogenetic trees from non-overlapping 5 kb windows and estimated a species tree from the 12,091 resulting window trees with ASTRAL v5.7.7 (Zhang *et al*., 2017) and default settings. Gene concordance factors (gCF) were calculated for the maximum likelihood and ASTRAL trees with IQ-TREE (**Methods S1)**. We also investigated phylogenetic relationships of the chloroplast and mitochondrial genomes of these samples (**Methods S1**).

### Comparative genomics

Syntenic structure for eight oak genomes, *Castanea mollissima*, and *Castanopsis tibetana* was assessed using SyRI as described above, with hapA as the reference **(Methods S1)**. A syntenic block analysis was conducted with 11 species of Fagales and *Prunus persica* (Rosales) by using OrthoFinder v2.5.4 (Emms & Kelly, 2019) on primary proteins, followed by syntenic block identification and duplicate gene classification by MCScanX (Wang *et al*., 2012; **Methods S1)**. Gene families were determined in hapA and primary protein sets from seven *Quercus* species by running GeneSpace (Lovell *et al*., 2022), which uses OrthoFinder v2.5.4 and MCScanX. Expansion and contraction of gene families was determined with CAFE5 (Mendes *et al*., 2020) and the base model. A summary of the taxa included in each comparative genomic analysis is shown in **Figure S1**.

## Results

### Haplotype-resolved genome assembly

The reference genome for *Quercus alba* (tree MM1, see **Figure 1**) was assembled with PacBio HiFi (circular consensus sequencing) reads with 45X haploid genome coverage and an average read length of 20,753 bases (**Table S2**). Assembly and scaffolding were further improved by a Hi-C library with 15X haploid genome coverage, which resulted in two resolved haplotypes (i.e. hapA and hapB). HapA includes 763 contigs spanning 794,299,596 bases (N50:29; L50:8.3 Mb), and 300 of these contigs, spanning 97.0% of the total bases, are scaffolded into 12 chromosomes (**Table 1**). HapB is similarly complete with 563 contigs (N50:29; L50:8.9 Mb) spanning 792,297,883 bases with 215 contigs representing 97.0% of the total bases scaffolded into 12 chromosomes. BUSCO analysis of the unannotated genomes using 1,614 Embryophyta conserved genes found that over 98% were present and complete in both haplotypes, and only 5% were complete and duplicated. A complete chloroplast and a draft mitochondrial genome in two scaffolds were also assembled for MM1 (**Figures S2-S3**).

**Table 1.** Assembly and BUSCO statistics from hapA and hapB, assessed for the total assembly (All) and the scaffolds placed into the 12 chromosomes (Chrs).

|  |  | HapA All | HapA Chrs | HapB All | HapB Chrs |
| --- | --- | --- | --- | --- | --- |
| Contigs | Number | 763 | 300 | 563 | 215 |
|  | Total bases | 794,299,596 | 770,521,868 | 792,297,883 | 768,546,887 |
|  | L50 | 29 | 28 | 29 | 28 |
|  | N50 | 8.3Mb | 8.4Mb | 8.9Mb | 8.9Mb |
| Scaffolds | Number | 417 | 12 | 309 | 12 |
|  | Total bases | 794,470,984 | 770,665,549 | 792,424,023 | 768,647,478 |
| Protein<br>Coding<br>Genes | Number | 42,955 | 42,489 | 42,412 | 42,150 |
| BUSCOS | Complete (% of 1,614 total<br>BUSCOs) | 1,592<br>(98.6%) | 1,579 (97.8%) | 1,591 (98.6%) | 1,587 (98.3%) |
|  | Complete and single-copy | 1,517 | 1,505 | 1,526 | 1,522 |
|  | Complete and duplicated | 75 | 74 | 65 | 65 |
|  | Fragmented | 9 | 13 | 14 | 14 |
|  | Missing | 10 | 22 | 9 | 13 |

A genetic map was constructed from the genotypes of 184 F1 progeny derived from an open-pollinated mother tree (WO1; **Figure S4; Table S3**) unrelated to the reference tree MM1. A total of 181 SNPs were organized into 12 linkage groups covering a total genetic distance of 731.5 cM, which is comparable with other *Quercus* genetic maps, including the 734 cM genetic map from *Q. robur,* the 764 cM genetic map from *Q. petraea*, and the 652 cM genetic map from *Q. rubra* (Bodénès *et al*., 2016; Konar *et al*., 2017; **Table S4**). The longest and shortest linkage groups were LG2 (89.2 cM) and LG9 (36.7 cM), respectively. Of the 177 genetic map markers with available sequence data, 167 and 166 mapped uniquely to the hapA and hapB assemblies, respectively (**Figure 2**). All markers from each linkage group mapped to the corresponding chromosome in each haplotypic assembly. Most markers were in the same order in the chromosomes as in the linkage map. Markers occurring in a different order on the genetic map compared to the genome assembly may be explained by a multigene family origin of markers or misplaced markers on the genetic map due to a relatively low number of F1 progeny.

**Figure 2.**
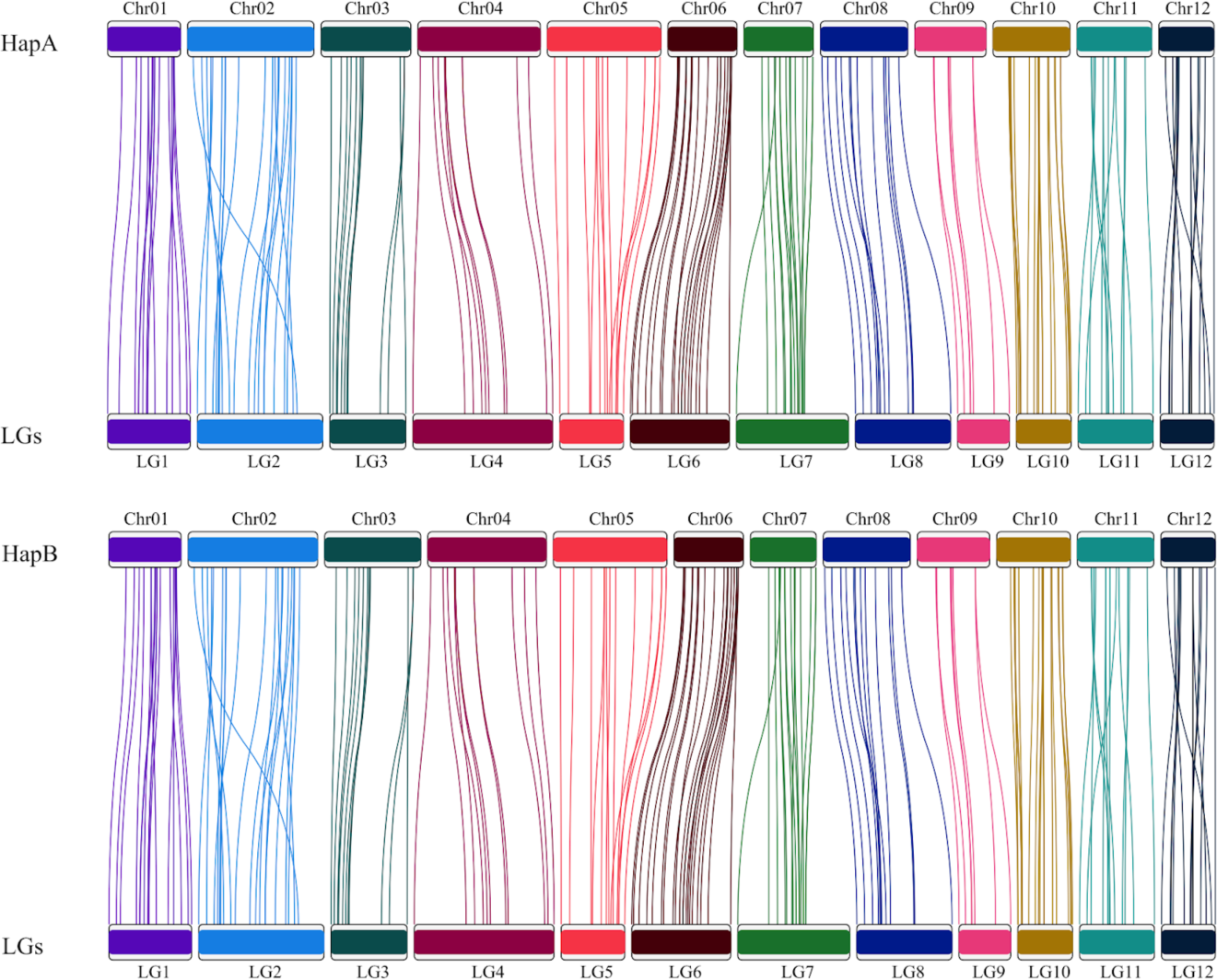
Comparison of a genetic map to the *Q. alba* genome assembly. The majority of markers from the genetic map (bottom) map uniquely to the chromosomes in hapA and hapB assemblies (top), visualized by colored lines.

HapA and hapB were highly collinear across all 12 chromosomes. Based on sequence alignment and analysis by SyRI, 1,283 syntenic blocks were identified, spanning 610 Mb in hapA and 620 Mb in hapB (**Figure 3**). Based on these alignments, SyRI (Goel *et al*., 2019) identified 12,808 structural variants (SVs) of at least 100 bases (**Table 2**). The majority of SVs involved fewer than 10,000 bases, and only two SVs were over 1 Mb in length: a 1.1 Mb inversion on Chromosome 3 and a 1.9 Mb inversion on Chromosome 9. An additional 24 SVs were over 100 kb in length, including 14 inversions, five translocations, and one 155 kb section on Chromosome 1 which was absent in hapA.

**Figure 3.**
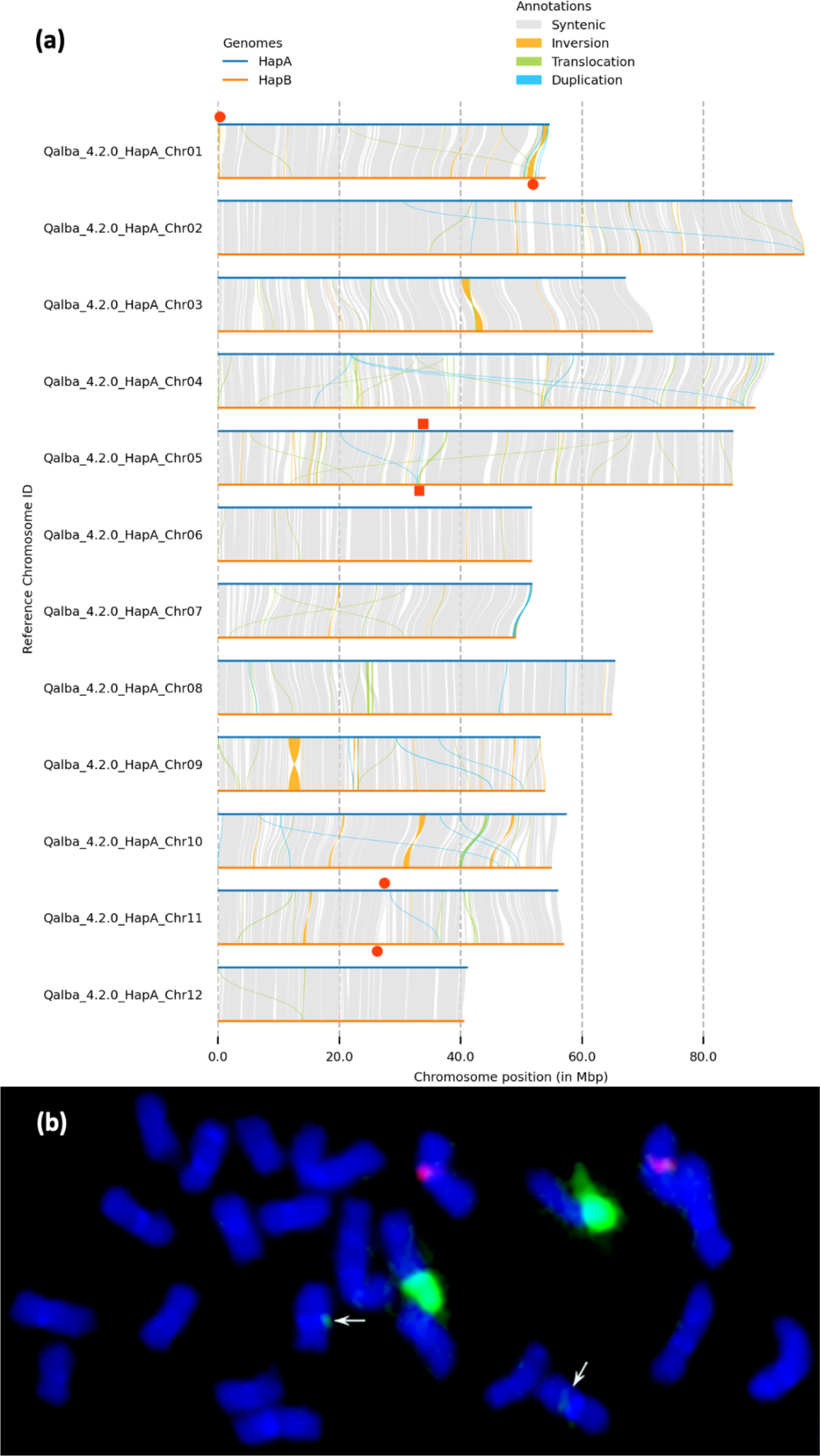
(a) Structural synteny between hapA and hapB of the *Q. alba* genome assembly. Two inversions are over 1 Mb: a 1.1 Mb inversion on Chromosome 3 and a 1.9 Mb inversion on Chromosome 9. The location of the 35S array is denoted by red squares, and the 5S arrays are denoted by red circles. **(b)** A metaphase chromosome spread with two pairs of 35S (green) and one pair of 5S (red) rDNA signals. Minor 35S signals are indicated by white arrows.

**Table 2.** Structural variation identified between the two assembled haplotypes of *Q. alba*.

|  | Total Count | 100 to < 1kb | 1kb to < 10kb | 10kb to < 100kb | 100kb to < 1Mb | > 1Mb |
| --- | --- | --- | --- | --- | --- | --- |
| Inversions | 103 | 40 | 18 | 29 | 14 | 2 |
| Translocations | 1,331 | 241 | 912 | 173 | 5 | 0 |
| Insertions | 4760 | 3621 | 1093 | 46 | 0 | 0 |
| Deletions | 4858 | 3730 | 1077 | 51 | 0 | 0 |
| Duplicated Regions | 803 | 170 | 595 | 38 | 0 | 0 |
| Inverted Duplicates | 652 | 138 | 431 | 83 | 0 | 0 |
| Tandem Repeat | 15 | 12 | 3 | 0 | 0 | 0 |
| Copy Gain in hapB | 134 | 61 | 54 | 16 | 3 | 0 |
| Copy Lost in hapB | 152 | 84 | 49 | 18 | 1 | 0 |
| ALL | 12808 | 8097 | 4232 | 454 | 23 | 2 |

### Structural and functional gene annotation

Repeats composed 58% and 59% of the hapA and hapB assemblies, respectively, with long terminal repeats (LTRs) as the largest class of elements at approximately 28% of the genome in both haplotypes (**Table S5**). The unplaced scaffolds had a higher percentage of bases identified as repetitive than the chromosomes: 67% vs 57% for hapA and 82% versus 58% for hapB, suggesting that they were not placed in a unique chromosome location likely due to the larger amount of repeat sequences interfering with the assembly process (Tørresen *et al*., 2019; Giani *et al*., 2020). The LAI (LTR Assembly Index), a metric of assembly quality based on completeness of LTRs (Ou *et al*., 2018), was 23.3 for hapA and 22.9 for hapB. These scores are in the top 10% of LAI scores of 103 high-quality plant genomes, indicating high assembly quality through repetitive regions for both white oak haplotypes. Categories of repeat types found in hapA and hapB were similar, except that the DNA transposon Tc1/mariner superfamily was found to constitute only 0.08% of hapA but made up 1.43% of hapB (**Notes S1; Table S5; Table S6**).

Protein-coding gene annotation identified 42,955 genes in hapA and 42,412 genes in hapB. Over 99% of genes were on chromosome scaffolds. Based on structural variants identified between the haplotypes, we found that 2,365 gene bodies (5.5% of genes) overlapped with an SV, and of those, 1,374 (3% of genes) had an exonic sequence overlapped by an SV. The most common type of SV to overlap a gene was an inversion (803 genes) with insertions as the second most prevalent type (554 genes).

Functional annotations were found for 82% of genes through sequence similarity to databases of proteins, metabolic pathways, and gene families (Zenodo accession TBD). Sixteen samples of RNA from four different white oak trees, including four emerging leaf samples from the MM1 tree, were used for annotation and then remapped to the annotated genome. Across samples, 81 to 91% of reads mapped uniquely, while 3 to 5% mapped to multiple locations. There were 29,594 genes in hapA that had at least one mapped RNASeq read and 23,639 had a TPM (transcript reads per kilobase of gene space per million) greater than 0.5. Tissues differed in number of genes expressed, from 17,582 genes in emerging leaves to 24,798 genes in the emerging radical apex of a germinating acorn (**Table S7**).

rDNA gene arrays were annotated in the genome by sequence similarity (**Figure 3a**). A single 5S array was found on chromosome 5 at 33.8 Mb in hapA (32.9 Mb in hapB). Two 35S arrays were found. One 35S array was located on chromosome 1 at 160 kb in HapA, but found at the opposite end of chromosome 1 in hapB (50-53 Mb). The second 35S array was found on chromosome 11 at 27.4 Mb in hapA (26.2 Mb in hapB). To confirm rDNA arrays, FISH was conducted and revealed one 5S locus and one 35S locus consistently, with an additional, possibly minor, 35S locus observed rarely, (i.e., not in every metaphase; **Figure 3b; Figure S5**). These sites are located on three different chromosome pairs, which is in agreement with the sequence-based analysis. The 5S and 35S sites appeared to be colocalized with AT-rich heterochromatic bands (**Figure S5**).

An assessment of annotated genes with R gene associated domains (CC, coiled coil; NBS, nucleotide binding region; LRR, leucine rich region; TM, transmembrane; TIR, Toll-interleukin region; or kinase) yielded 2,503 genes in hapA and 2,292 genes in hapB (**Table 3**). Based on the domains, the genes were further classified into four main categories (**Table 3**). RLP (receptor-like proteins) were the most common, followed by RLK (receptor-like kinases), CNL (CC-NB-LRR), and TNL (TIR-NB-LRR) as the smallest gene category.

**Table 3.** R gene domains and R gene classifications identified between the two assembly haplotypes of *Q. alba*.

|  | Genes in hapA | Genes in hapB |
| --- | --- | --- |
| **Domains** |  |  |
| Transmembrane Helix | 2093 | 1922 |
| Kinase | 1498 | 1426 |
| Leucine-rich Repeats | 1090 | 945 |
| Nucleotide-binding Site | 467 | 410 |
| Coiled-coil | 331 | 286 |
| Toll/interleukin-1 Receptor | 49 | 37 |
| **R Gene Classifications** |  |  |
| Receptor Like Proteins | 381 | 355 |
| Receptor Like Kinase | 302 | 285 |
| CC-NB-LRR | 102 | 81 |
| TIR-NB-LRR | 19 | 11 |

### Clustering, principal component analysis, and population genetics

In addition to our reference individual, we performed whole genome shotgun sequencing of 18 *Q. alba* individuals representing much of the species rangewide diversity (**Table S1**). Our filtered dataset for the 19 *Q. alba* samples consisted of 50,461,765 SNPs, 7.5% of which had three or more alleles (including single nucleotide deletions). We summarized our *Structure* results with CLUMPAK, which revealed high consistency among replicate runs for values of *K*= 1, 2 and 3, with all 10 replicates clustering into a single mode per value of *K* (**Figure S6**). PCA analysis revealed that the first two principal component axes strongly correlated with the geographic origin of samples (**Figure 4**). The clustering pattern of individuals in the PCA was also qualitatively consistent with *Structure* results at *K*=3, and we therefore used this value of *K* to define groups for our “genetic clustering” set of population genetic analyses.

**Figure 4.**
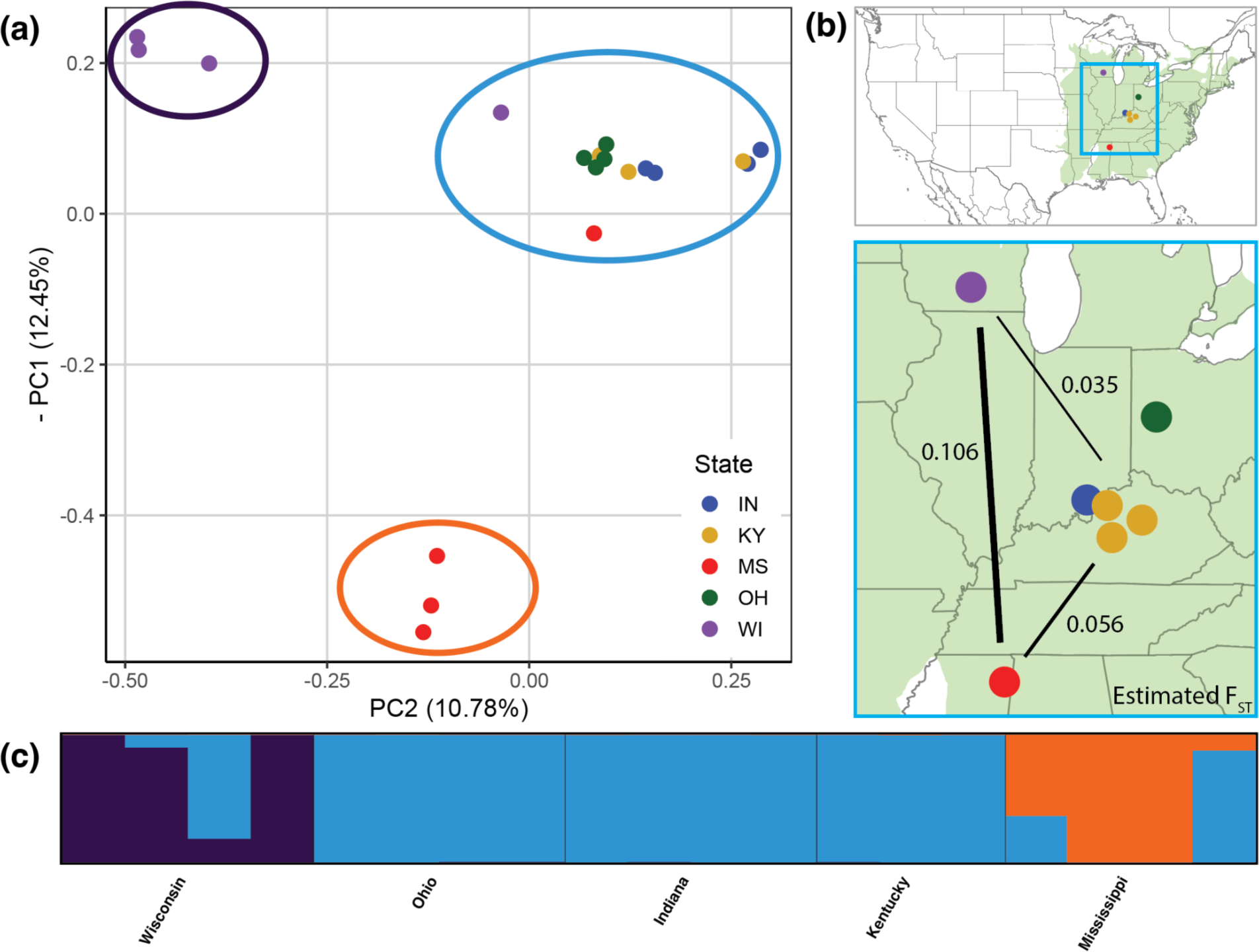
Population structure within *Q. alba*. A) Principal component analysis of 19 *Q. alba* individuals. The inverse of PC 1 is plotted on the y-axis and PC 2 is plotted on the x-axis. Colored circles around clusters correspond to the color scheme of panel C and denote the major ancestry of those individuals in *Structure* analyses for *K*=3. B) Map of sample provenances colored by state with the range of *Q. alba* shown in green. Numbers and black line thickness correspond to *F*ST (Weir and Cockerham estimator) between groups of samples from Wisconsin, Mississippi and a combined group from Indiana, Ohio, and Kentucky. C) *Structure* result for a typical replicate with *K*=3 for the 19 individuals of *Q. alba* from five US states.

When grouping individuals by state, pairwise *F*ST with the Weir and Cockerham (1984) estimator was greatest between samples from Wisconsin and Mississippi (*F*ST = 0.106; **Table S8**). For “geographically clustered” analyses, estimated *F*ST between the Ohio, Indiana, and Kentucky group and either the Wisconsin or Mississippi groups was 0.035 and 0.055, respectively (**Figure 4C**). For “genetically clustered” analyses of *F*ST, the estimated values for Northern vs. Southern, Northern vs. Central, and Central vs. Southern were 0.175, 0.049, and 0.082, respectively. Values of *F*ST with the Hudson *et al*. (1992) estimator were similar in their relative magnitudes, but were consistently higher than with the Weir and Cockerham estimator **(Table S8)**.

The value of π (nucleotide diversity) was greatest when considering all 19 *Q. alba* individuals as a single population (**Table 4**). When grouping individuals by state of origin, the Kentucky group had the largest π, followed by those from Indiana, Wisconsin, and Mississippi. When grouping by “geographic clusters,” the group consisting of samples from Ohio, Indiana, and Kentucky had the largest π. Similarly, the central cluster had the largest π in the “clustered individuals analysis” (**Table 4**).

**Table 4.** Estimated nucleotide diversity (π) for 19 *Q. alba* samples analyzed as a single group, grouped by state, clustered geographically (samples from OH, IN, and KY grouped together), or clustered genetically (based on major ancestry in *Structure* for *K*=3).

| Analysis | Group | $\pi$ |
| --- | --- | --- |
| All samples | - | 0.0120 |
| By state | Kentucky | 0.0117 |
|  | Indiana | 0.0107 |
|  | Wisconsin | 0.0102 |
|  | Mississippi | 0.0099 |
|  | Ohio | 0.0095 |
| Clustered geographically | OH+IN+KY | 0.0118 |
|  | Wisconsin | 0.0102 |
|  | Mississippi | 0.0099 |
| Clustered genetically<br>(*structure*) | Central | 0.0118 |
|  | Northern | 0.0095 |
|  | Southern | 0.0093 |

### Phylogenomic analyses

We produced whole genome shotgun sequencing reads for individuals of seven North American white oak species, which were analyzed along with public short-read data from an additional 15 *Quercus* species and *Lithocarpus longipedicellatus* **(Table S1)**. Our final phylogenomic dataset consisted of 327,242,758 aligned sites that were not masked in all samples. Based on our maximum likelihood tree, all individuals of *Q. alba* formed a clade that was sister to *Q. montana* **(Figure S7)**. All individuals from North American white oak species formed a clade, which was sister to a clade composed of *Q. robur*, *Q. petraea*, and *Q. mongolica*, the other three white oak species in our sampling (**Figure 5**). Section *Lobatae*, represented by *Q. rubra*, was sister to the white oak clade.

**Figure 5.**
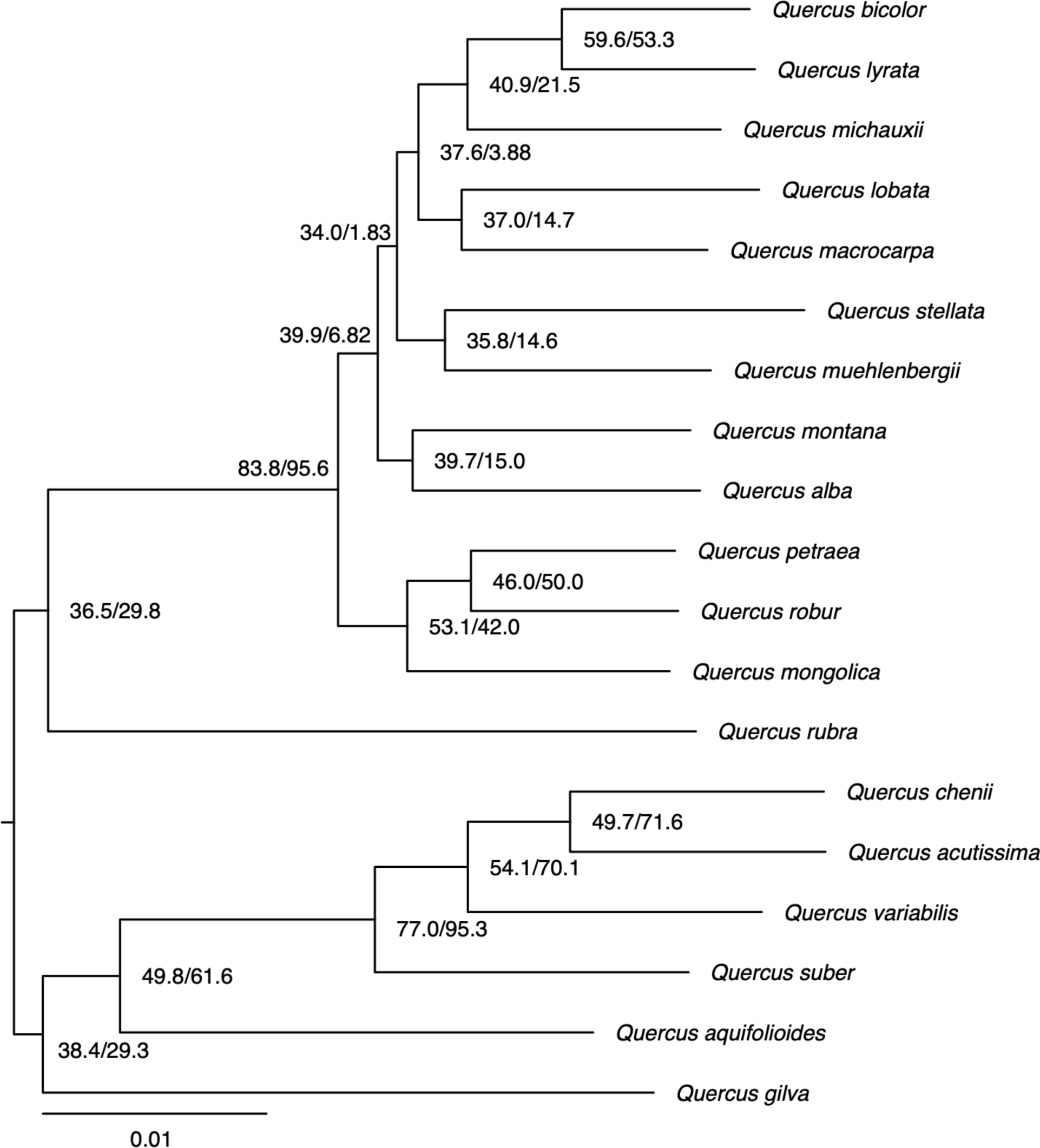
Phylogenetic tree estimated with maximum likelihood from a concatenated alignment of genome sequences from 19 oak species and rooted on an individual of *Lithocarpus* (not depicted). Node values are site concordance factors (left) and gene concordance factors (right) All species relationships received 100% ultrafast bootstrap support. Branch lengths are in units of estimated substitutions per site.

Both site concordance factor (sCF) and gene concordance factor (gCF) values indicated extensive phylogenomic discordance for many relationships in the tree, especially within the more intensively sampled white oak clade. Within this clade there are several branches that were supported by fewer than 10% of genomic windows. There was no tree for any 5-kb window that was completely concordant with the inferred species trees. The ASTRAL species tree was similar in topology to the maximum likelihood tree, except for several relationships within the white oak clade, which were subtended by extremely short branches **(Figure S8)**. In the chloroplast and mitochondrial trees, *Q. alba* was not monophyletic and both trees were characterized by short internal branches within section *Quercus* and by widespread phylogenetic conflict with one another (**Figures S9-S11**).

There were 4,417,437 sites (singletons excluded) inferred to be variable among *Q. alba* individuals; of these, 57.7% of sites were inferred to be variable among other white oak species (**Figure S7**, **Table S9**). When singletons were included, there were 16,388,119 variable sites within *Q. alba*, with 43.6% of those also variable among other white oak species. Correcting for this ancestral variation resulted in divergence time estimates that were up to 9.9 million years closer to present than analyses that used uncorrected branch lengths **(Figure S12)**.

### Genomic architecture across the Quercus clade

Of the *Quercus* species included in our phylogenetic analysis, eight in addition to *Q. alba* have available chromosome-scale genomes, providing an opportunity to examine the evolution of chromosome structure across the genus (**Figure S1, Table S10**). Using *Q. alba* as the reference, the genomes were aligned and assessed for structural variation, revealing that they share strong chromosome-to-chromosome syntenic structure (**Figure 6A, Figure S13**). Within homologous chromosomes, numerous small- and medium-sized structural variations were identified between *Q. alba* and the other *Quercus* species (**Table S10**). There did not appear to be a clear relationship between structural variation and phylogenetic relatedness to *Q. alba*; the total number of structural variants found varied from 4,631 with *Q. variabilis* to 11,324 with *Q. lobata*. The largest structural variant in each comparison varied from a 1.9 Mb inversion in comparison to *Q. robur* to an 11.4 Mb inversion in comparison to *Q. acutissima* (**Table S10**).

**Figure 6.**
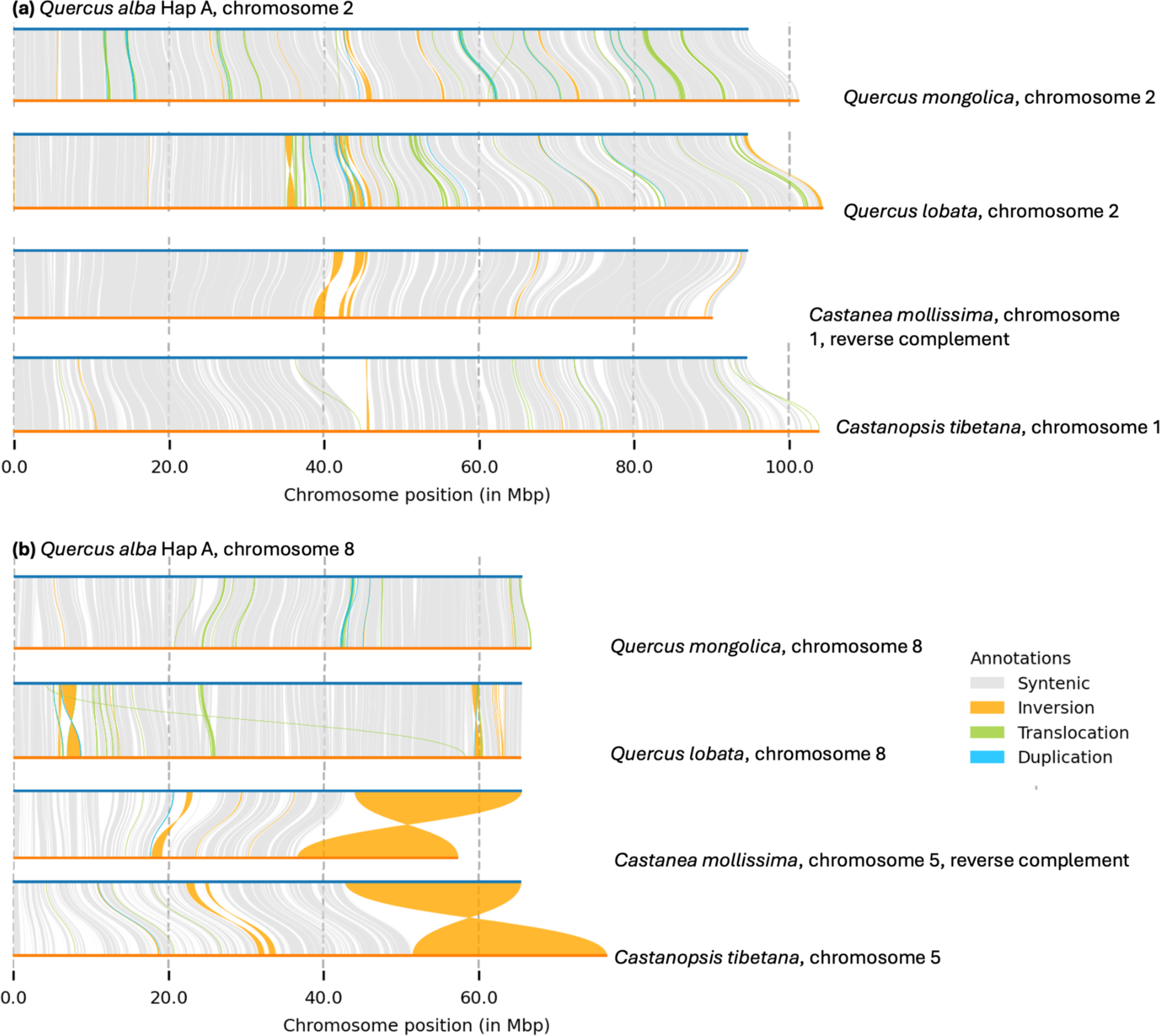
Structural synteny of *Q. alba* hapA (top bars, blue) versus homologous chromosomes (bottom bars, orange) of *Q. mongolica*, *Q. lobata*, *C. mollissima*, and *Castanea tibetana*. **(a)** An example of a typical comparison (chromosome 2), showing strong collinearity overall with small structural rearrangements. **(b)** Chromosome 8, for which investigated members of *Quercus* share a large inversion relative to other Fagaceae. Comparisons of the remaining chromosomes across the same four species are available in Supplemental Figure 3.

Two species from other genera within Fagaceae with chromosome-scale genomes, *Castanea mollissima* and *Castanopsis tibetana,* were also examined for syntenic structure. A major structural variant, a 22 Mb inversion at the end of Chromosome 8, was found to be nearly identically located in both non-oak species (**Figure 6B**) but was not found in any *Quercus* species. Further manual curation of structural variants over 1 Mb identified three additional smaller inversions (2.1 Mb on Chr 1, 1.1 Mb on Chr 7, and 1.0 Mb on Chr 8) that appear to be in a single orientation shared by all *Quercus* species and in the opposite orientation in the two non-oak species (**Figure S14**).

Examining species outside Fagaceae, many major chromosomal alterations are evident, and chromosomal segments that can be directly mapped based on sequence similarity are shorter. However, despite nucleotide divergence, gene collinearity is often conserved in large blocks (**Figure 7A; Figure S1**). To further characterize the nature of gene duplication across this same set of tree species, we used MCScanX to classify genes as originating from whole genome duplications, tandem duplications, proximal duplications, dispersed duplications, or single copy genes (i.e. no duplication). Interestingly, all species in the Fagales share very similar relative percentages of these categories, with most genes classified in the dispersed duplicates category (**Figure 7B**).

**Figure 7.**
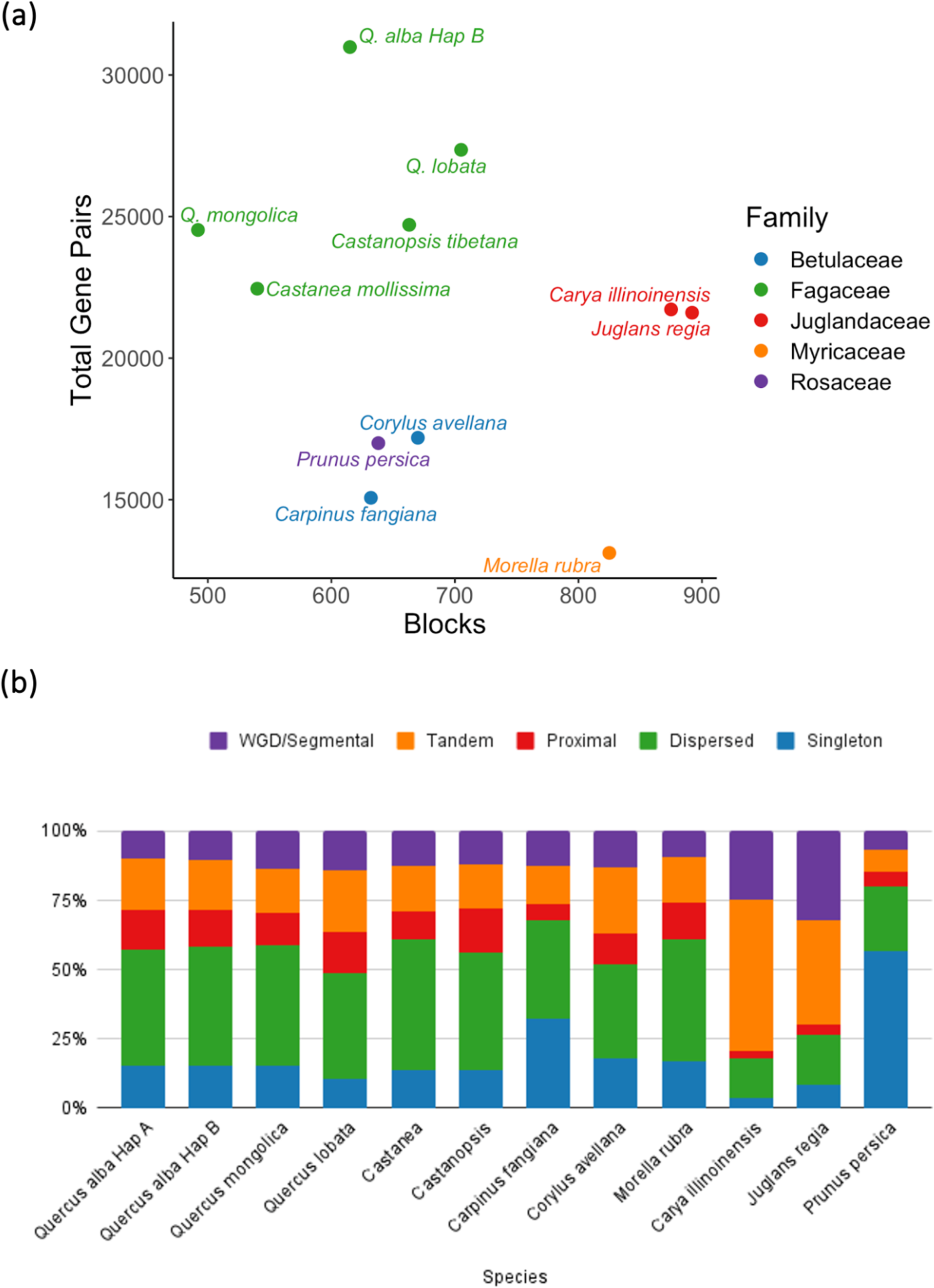
Comparative analysis of genome collinearity and gene duplications between *Q. alba* and tree species from four additional rosid families. **(a)** Comparison between the number of blocks of collinear genes and the total number of collinear genes identified. **(b)** Relative percentage of gene pairs within a genome generated by whole genome duplications (WGD), tandem duplications, proximal duplications, dispersed duplications, or no duplication, i.e. singleton genes.

### Gene family evolution among Quercus

The combination of fully annotated genomes and an ultrametric phylogeny of *Quercus* enabled examination of the evolution of gene families across the genus. Using all annotated protein-coding genes with chromosomal placement from seven *Quercus* species, Orthofinder identified 30,270 total orthogroups (mean size of 7 genes) and 13,623 orthogroups with all species present. CAFE was used to model gene family evolution and to detect families with significantly accelerated gene gain or loss on each branch of the phylogenetic tree. CAFE identified 852 gene families in *Q. alba* that have rapidly evolved in gene copy-number since the last common ancestor with *Q. lobata* (p < 0.05), reflecting a gain of 2,355 genes and a loss of 611 genes (**Figure 8A**). Based on gene ontology enrichment analysis (p < 0.05) of these rapidly evolving families, of the top twenty significantly enriched biological process terms, seven of those terms related to external stimulus and stress response (**Table S11**). Two additional terms relate to growth regulation. We identified 89 significantly changing gene families along the branch subtending section *Quercus*, with 269 genes gained and 25 genes lost (**Figure 8B**). Of the top 20 GO terms enriched in these rapidly evolving families, six were related to defense (**Table S12**).

**Figure 8.**
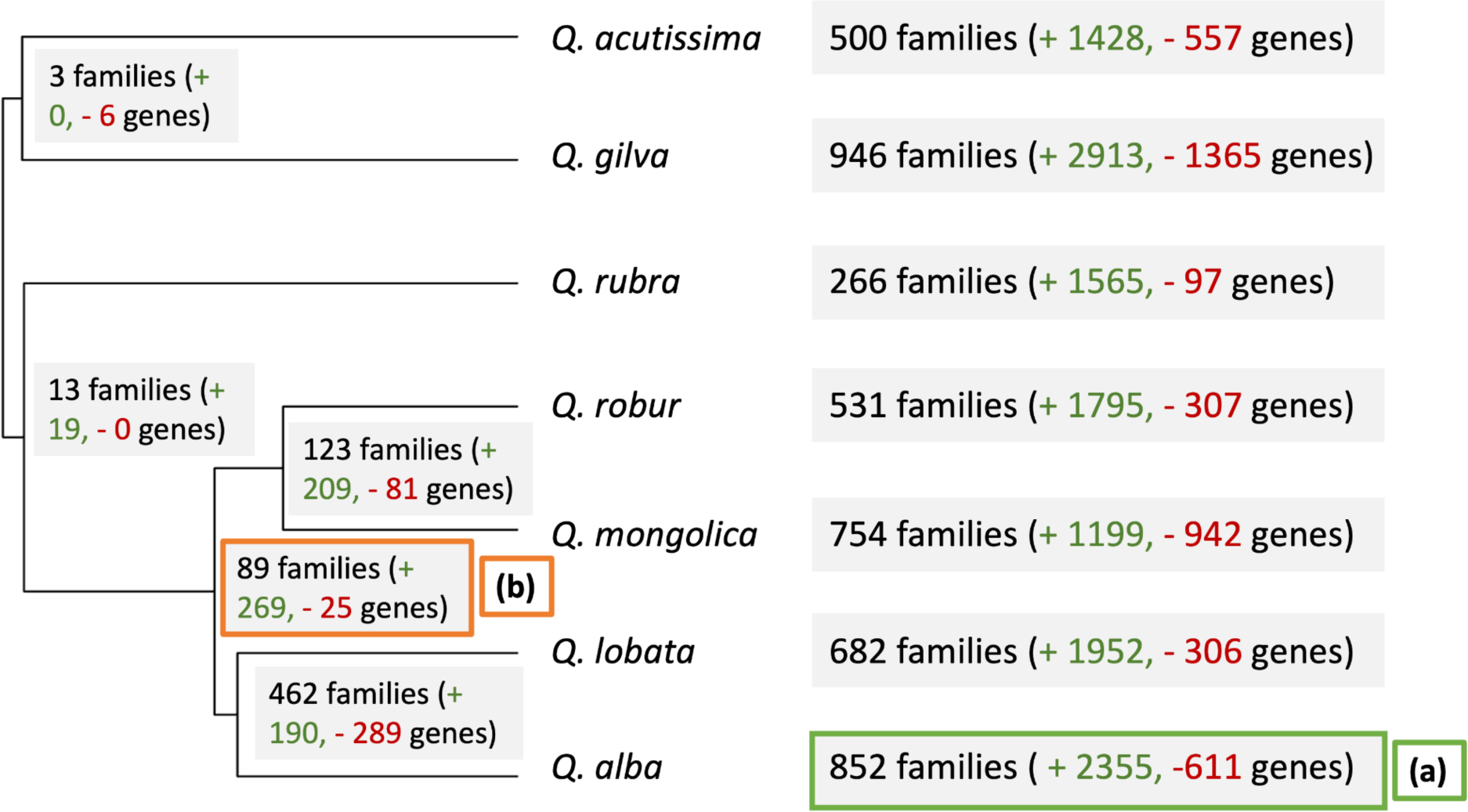
CAFE detected gene families with significantly accelerated gene gain or loss on each branch of the phylogenetic tree (grey boxes). Gene ontology enrichment was conducted to functionally profile the rapidly changing gene families in the *Q. alba* genome since its most recent ancestor with *Q. lobata* **(a)** and for the white oak clade (section *Quercus*) after branching from the rest of the *Quercus* genus **(b)**.

To further examine the evolution of defense response genes, we ran the same R gene identification pipeline used for *Q. alba* on seven annotated *Quercus* genomes (**Figure 9A**). The number of R genes overall and the number of genes per category (RLK, RLP, CNL, and TNL) varied widely among the species; however, RLKs and RLPs were more common than CNL/TNL in all species. The TNL gene family ranged from no identified members in *Q. gilva* to 241 genes in *Q. lobata*. We identified orthogroups composed primarily of R genes and found that these overlapped with many of the orthogroups with significant changes identified with CAFE, suggesting that some R gene families are rapidly evolving. All four R gene categories had expansions and contractions across the *Quercus* phylogeny. The mechanisms of gene expansion and contraction were identified using a duplicate gene classifier tool (**Figure 9B, Figure S15**). Examining all genes in the genomes, all species other than *Q. rubra* have a majority of genes originating from dispersed duplication. R genes are not reflective of the same overall gene duplication pattern: in contrast to the entire gene set, R genes are more likely to originate from proximal and tandem repeats. No RLKs, CNLs or TNLs in any species were identified as singletons.

**Figure 9.**
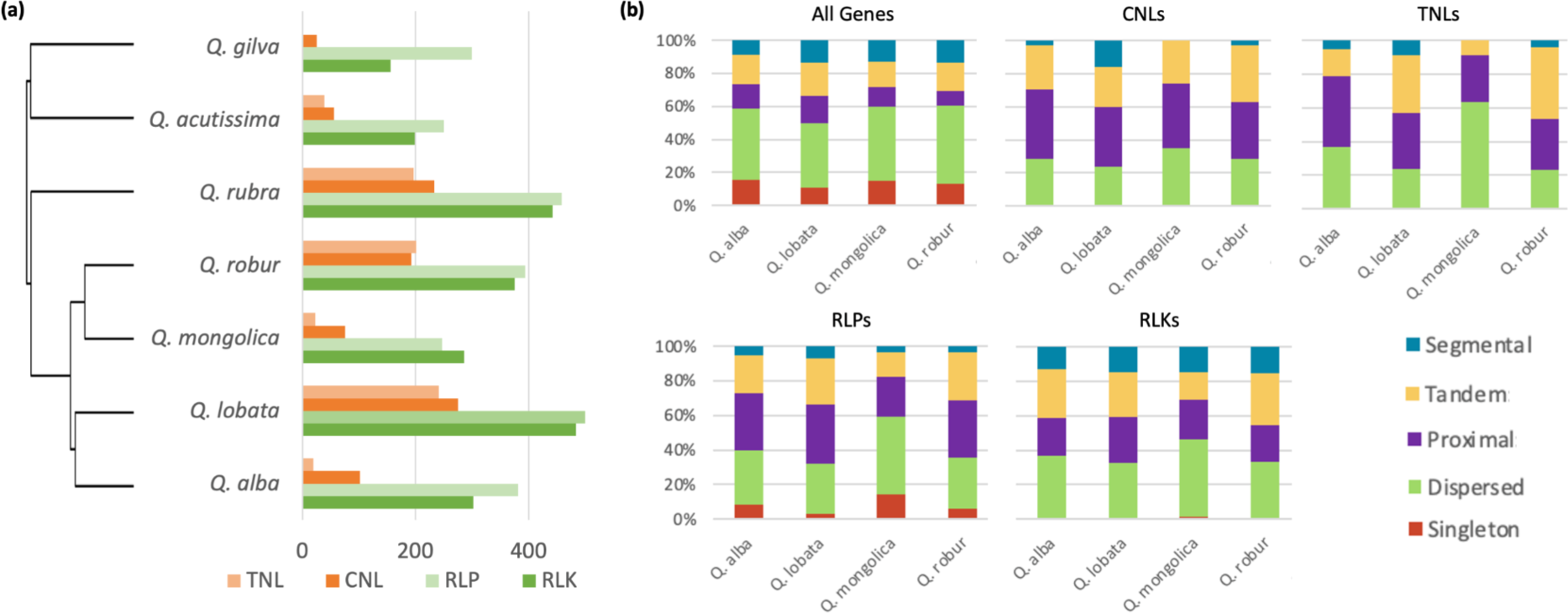
(a) The number of R genes that were recovered from *Quercus* genomes in four main categories: TNL (TIR-NB-LRR), CNL (CC-NB-LRR), RLP (receptor-like proteins), and RLK (receptor-like kinases). **(b)** Gene duplicate classification assigned genes to their most like origin: segmental duplication (gene is in a collinear block with homology to another colinear block), tandem (gene has homology to a neighboring gene), proximal (gene has homology to a gene within 20 genes upstream or downstream on the chromosome), dispersed (gene has homology to a gene somewhere else in the genome), or singleton (no detectable history of duplication). Analysis was conducted for all genes in each genome and for each of the four categories of R genes: CNLs, TNLs, RLPs, and RLKs. Patterns for *Q. alba* and three additional *Quercus* species are shown; analyses of additional *Quercus* genomes are in **Figure S15**.

## Discussion

### The Quercus alba genome

We generated a haplotype-resolved and chromosome-scale *Q. alba* reference genome that will support new genetic and genomic research in forest trees, an area of plant genomics neglected in comparison to crop plants (Neale & Kremer, 2011; Plomion *et al*., 2016b). Genomics can be a key component of creating a sustainable supply of white oaks for ecosystem management, forest restoration, and wood products such as lumber, veneer, and barrels for aging wine and spirits (Grattapaglia *et al*., 2009; Wheeler *et al*., 2015). There are currently active tree improvement programs conducting progeny trials and establishing seed orchards to supply genetically improved white oak acorns for nursery production (Schlarbaum, 1993, 2000, 2024; Dewald *et al*., 2023). These programs currently select for heritable traits such as rapid early growth and apical dominant architecture, but are limited in progress by multi-decade generation times and huge space requirements. Leveraging the *Q. alba* reference genome to map traits and to establish molecular screening approaches for early selection of high performing material or to implement a breeding program with genomic selection promises to rapidly advance the quantity and quality of white oak germplasm (Wheeler *et al*., 2015). The white oak genome will also open avenues to study genes underpinning traits characterized in other tree species, including wood-quality traits such as lignocellulose composition and optimization of flavor expression in oak barrels used for spirit aging (Grattapaglia *et al*., 2018; Gollihue *et al*., 2018, 2021).

We confirmed the completeness and accuracy of both genome assembly haplotypes with BUSCO and LAI scores as well as a new genetic linkage map. The haplotypes have contig N50s of 8.3 Mb and 8.9 Mb, commensurate with the highest quality recent plant genome assemblies (Kong *et al*., 2023), with 97% of bases scaffolded into chromosomes (**Table 1**). Similar to other *Quercus* genomes (Sork *et al*., 2022; Ai *et al*., 2022), each haplotype has 12 chromosomes and is approximately 793 Mb in length, in agreement with cytology (Friesner, 1930) and C-value estimates (Bai *et al*., 2012). The availability of both assembled haplotypes created an opportunity to explore structural variation (SV) within a single individual’s genome. The haplotypes have highly similar overall structures as expected, but also extensive structural variation (**Figure 3**). We identified over 12,000 structural variants, with insertions and deletions as the most common type, followed by duplications and translocations (**Table 2**). SVs are generally deleterious, but have also been identified as drivers of adaptation and the causative mutations underlying phenotypic differences in woody plants (Zhou *et al*., 2019; Guo *et al*., 2020; Hämälä *et al*., 2021).

Supporting their potential importance in the white oak genome, we found that 5.5% of gene bodies overlapped SVs and 3% of genes overlap SVs in exonic regions. Indeed, further range wide evaluation of SVs in *Q. alba* is needed to carefully evaluate their adaptive importance.

It is still unclear if the large structural variants we observed originated from assembly or scaffolding errors, particularly for SVs that are longer than individual PacBio HiFi reads (Yuan *et al*., 2021). While the single 5S array and one of two 35S arrays were placed in the same chromosomal locations between hapA and hapB, the other 35S array was placed on opposite ends of chromosome 1 (**Figure 3**). rDNA arrays have been found particularly difficult to assemble in other species as well (Navrátilová *et al*., 2022; Huff *et al*., 2023) and we posit that due to the location in the highly repetitive telomeric region of chromosome 1, our methods were unable to consistently place the 35S array. The confirmation of large structural variants, particularly those in repetitive regions, will require additional investigation.

### The evolution of gene family size across the *Quercus* phylogeny

Disease resistance genes (R genes), also referred to as pathogen recognition genes (PRGs) encompass multiple categories of defense genes with conserved domain patterns essential to pathogen recognition and defense initiation, with many R gene types ubiquitous across land plants (Yue *et al*., 2012; Fischer *et al*., 2016; Liu *et al*., 2017). Genes functioning in defense have been identified as some of the most rapidly evolving gene families in plants with signatures of both purifying and positive selection (Fischer *et al*., 2016; Zheng *et al*., 2016). We analyzed rapidly changing gene families along two branches of our phylogenetic tree: the terminal branch leading to *Q. alba*, and the branch subtending section *Quercus*. In both cases, defense related genes were strongly enriched, suggesting that, despite long generation times, R gene families appear to be capable of rapid evolution in tree species. It should be noted that CAFE is not able to model gene tree discordance, which may lead to increased false positive rates of rapidly evolving families (Neafsey *et al*., 2015; Mendes & Hahn, 2016).

After the initial assembly of the first oak reference genome, *Q. robur*, Plomion et al. noted an enrichment of R genes compared to other plant species (Plomion et al., 2018). This was echoed by Sork et al., who further noted the mechanisms of tandem duplications for R gene proliferation in *Q. lobata* (Sork *et al*., 2022). In contrast, Ai *et al*. (2022) identified a noted decrease in R genes in *Q. mongolica* (Ai *et al*., 2022). We analyzed four specific R gene categories in *Q. alba* and seven additional *Quercus* species. We found that R genes are, similarly to patterns observed in most plant clades, rapidly evolving and vary in number throughout *Quercus*. This suggests that, at least within *Quercus*, long life span may not be directly correlated to R gene count. Furthermore, R gene content appears to be largely species specific rather than showing a clear phylogenetic pattern. Future work characterizing additional *Quercus* genomes will likely shed additional light on the evolution of the R genes in oaks. While analysis of both the *Q. lobata* genome (Sork *et al*., 2022) and the *Q. robur* genome (Plomion *et al*., 2018) highlighted tandem and proximal duplication as a central hallmark of R gene evolution, we found dispersed duplication is an equally or more common predicted mechanism for R gene expansion.

### Genomic diversity of *Quercus alba*

We found evidence that *Q. alba* maintains high genetic diversity across its range (**Figure 4**). Our overall estimate of nucleotide diversity (π) for *Q. alb*a was 1.2%, which is similar to estimates from other broadly distributed oak species (Plomion *et al*., 2018). There was no clear geographic trend in genetic diversity, though samples from Wisconsin and Mississippi, representing the most northern and southern populations, respectively, showed a value of π that was slightly lower than samples from Kentucky and Indiana (two states in the central part of *Q. alba*’s range). Our results also showed population structure within *Q. alba*; both principal component analysis and *Structure* analysis revealed genetic differentiation among samples. Samples from the northern and southern populations largely formed distinct clusters, whereas the remaining samples clustered together. This pattern of clustering is also evident in our phylogenetic results (**Figure S7**). While there is some evidence from provenance trials that some populations of white oak may be locally adapted, including exhibiting differences in leaf phenology (Huang *et al*., 2016), there remains much to be learned about genetic differentiation across the species range. Our results suggest that there is a clear potential for local adaptation in white oak due to the presence of population structure within the species. The *Q. alba* genome will be a useful tool for future studies aimed at characterizing the genomic basis of local adaptation in the white oak clade.

### Shared variation among oak species and implications for divergence time estimation in oaks

We found that there are millions of shared variable sites among white oak species, which has important implications for phylogenetic analysis and divergence time estimation. By some metrics, as many as 57.7% of variable sites within *Q. alba* were also variable among other white oak species (**Figure S7**). Such a result suggests that much of this variation has been shared since their common ancestor.

When analyzed in a typical phylogenetic framework, this shared variation is ignored. As a result, when different alleles are sampled in different individuals, these may appear as differences between individuals or species. This effect was apparent in our results: in many cases, the inferred divergence between individuals within *Q. alba* was nearly as high as between several species pairs (**Figure S7**). An important downstream effect of mistaking ancestral variation for differences that have accumulated since speciation is the inference of erroneously long branch lengths, which has the effect of “pushing back” estimated divergence times (Edwards & Beerli 2000). To account for this effect, we used our estimate of π from *Q. alba* to correct branch lengths and to account for this ancestral diversity, which for many nodes resulted in divergence times that were 8-10 million years nearer to the present than when using uncorrected branch lengths analyzed with the same dating methods **(Figure S12)**.

### The history of the white oak clade is characterized by extensive phylogenomic conflict

Our results using whole genome alignments suggest a different phylogenetic history for some taxa compared to a recent, broadly sampled phylogenetic analysis based on RAD-Seq data (Hipp *et al*., 2020), though our results agree for many relationships (Notes S2). Ubiquitous phylogenomic conflict, geographic variation in patterns of introgression, and sampling and sequencing differences could contribute to differences in our inferred species relationships compared to previous studies. While detailed investigation of patterns of hybridization and introgression are beyond the scope of this work, we found high phylogenetic discordance across the white oak clade and that a large proportion of sites that were variable within *Q. alba* were also variable among other species of white oaks (**Figure S6**). These findings could be due, in part, to introgression as well as incomplete lineage sorting following rapid radiations in the clade. Oaks are often reported to hybridize, e.g. (Leroy *et al*., 2017; Hipp *et al*., 2019; Degen *et al*., 2023). Hardin (1975) described morphological evidence of hybridization between *Q. alba* and nine other species in the white oak clade, which includes ca. 15 other species in eastern North America. Hybridization followed by backcrossing can facilitate the transfer of adaptive alleles across species boundaries, as has been suggested by recent evidence in European white oaks (Leroy *et al*., 2020). Future work should address the role of introgression in the white oak clade using whole genome data.

## Acknowledgements

Sincere thanks to Rob Samuels and Brian Mattingly of Maker’s Mark Distillery and Brad Boswell and the Boswell Family of the Independent Stave Company for their initiative, financial support and passion for advancing science. Support was also provided by the National Science Foundation: IOS-2109716 to DAL, DBI-2146866 to MWH, and IOS-1025974 to JEC, MES, and SES. Additional support came from McIntire-Stennis Project 4717 to JEC and from USDA Forest Service, Southern Research Station Research Joint Venture Agreement 19JV11330126084 with the University of Kentucky, College of Agriculture, Food and the Environment. Thanks to Mark Coggeshall (retired USDA Forest Service, Northern Research Station) and Phil O’Connor (Indiana Department of Natural Resources) for establishing and maintaining the “postage stamp” planting in Indiana where the provenanced *Q. alba* trees were sampled, Dr. Catherine Bodenes (INRA, France) for assistance in constructing the genetic linkage map, and to Jess Slade (The Arboretum at the University of Kentucky) for assistance in locating trees for sampling. This research was supported in part by an appointment to the United States Forest Service (USFS) Research Participation Program administered by the Oak Ridge Institute for Science and Education (ORISE) through an interagency agreement between the U.S. Department of Energy (DOE) and the U.S. Department of Agriculture (USDA). ORISE is managed by ORAU under DOE contract number DE-SC0014664. All opinions expressed in this paper are the authors’ and do not necessarily reflect the policies and views of USDA, DOE, or ORAU/ORISE.

## Author contributions

DAL, MES, SES, TZ, MWH, JEC, BA, SD, and DN designed the research. DAL, MES, BK, SF, JS, NF, SES, AH, ECS, TZ, JEC, and DN performed the research. DAL, MES, BK, SF, JS, AT, NF, ASA, ECS, TZ, MWH, JEC contributed to data analysis, collection, and/or interpretation. DAL and MES wrote the manuscript with contributions from BK and NF. All authors reviewed and approved the manuscript.

## Data availability

Raw sequences have been submitted to NCBI Sequence Read Archive and aggregated under BioProject PRJNA1021599. HapA is NCBI genome GCA_036321655.1 and HapB is NCBI genome GCA_036321645.1. Individual sample accession numbers can be found in the supporting information accompanying this article (Tables S1-S2). Novel scripts used in the analyses underlying this article, functional gene annotations for the white oak genome, and additional datasets and output are available from Zenodo: [Accession number to be determined]

## List of supporting materials

**Methods S1.** Supporting methods.

Table S1. Sample locations and NCBI data accessions for trees used for population genetics and phylogeny analysis.

Table S2. DNA and RNA sequencing data statistics

Table S3. Genetic Map Markers, Locations, and Sequences

Table S4. Genetic linkage maps of Quercus robur and Quercus petrea (Bodénès et al., 2016), Quercus rubra (Konar et al., 2017) and Quercus alba (this study).

Table S5. Repeat Profiles

Table S6. Tc1/Mariner Superfamily Profiles

Table S7. Genes expressed by tissue

Table S8. FST values with the estimators of Weir and Cockerham (1982) and Hudson et al. (1992) for individuals grouped by state, geographic clusters and genetic clusters.

Table S9. Shared variable sites analysis results.

Table S10. Structural variation of *Q. alba* vs 10 Fagales genomes.

Table S11. GO terms enriched in rapidly evolving gene families in *Q. alba* since its most recent common ancestor with *Q. lobata*

Table S12. GO terms enriched in rapidly evolving gene families in *Quercus* section *Quercus*.

Figure S1. Summary of taxa included in each comparative genomics analysis.

Figure S2. The white oak (MM1) chloroplast genome.

Figure S3. Annotated, circularized draft assembly of the *Quercus alba* (MM1) mitochondrial genome.

Figure S4. Female WO1 map composed of 181 SNP markers.

Figure S5. Fluorescence in situ hybridization (FISH) of white oak chromosome spreads reveals two pairs of 35S (green) and one pair of 5S (red) rDNA signals.

Figure S6. Major modes recovered with CLUMPAK for *Structure* results for K=1 to 5.

Figure S7. Phylogenetic tree of *Quercus* including all sampled individuals of *Quercus alba* and results regarding shared variable sites.

Figure S8. ASTRAL tree, generated from 12,081 gene trees based on 5 kb windows.

Figure S9. Chloroplast genome tree.

Figure S10. Mitochondrial genome tree.

Figure S11. Topological comparison between the plastome (left) and mitogenome (right) trees.

Figure S12. Dated phylogenies.

Figure S13. Overview of structural synteny of *Quercus* genomes, from top to bottom: *Q. rubra* (Qru), *Q. robur* (Qro), *Q. mongolica* (Qmo), *Q. lobata* (Qlo), *Q. gilva* (Qgi), *Q. alba* (Qal), *Q. acutissima* (Qac).

Figure S14. Detailed structural synteny of *Q. alba* Hap A against *Q. mongolica* (upper left), *Q. lobata* (upper right), *Castanea mollissima* (lower left), and *Castanopsis tibetana* (lower right).

Figure S15. Gene duplicate classification of all genes and R genes from *Quercus* genomes

