## Supplemental Materials for "A haplotype-resolved reference genome of *Quercus alba* sheds light on the evolutionary history of oaks"

### New Phytologist Supporting Information

**The following Supporting Information is available for this article:**

**Supporting Methods**

**Methods S1.** Supporting methods.

**Reference genome tissue collection, high molecular weight DNA extraction, and sequencing**

An individual of *Quercus alba* in a forest stand near Loretto, Kentucky, USA (37° 39.0583', -85° 21.2615'), referred to as MM1 (**Figure 1**), was sampled and vouchered with permission of the landowners. Emerging leaves were collected, flash frozen in liquid nitrogen, and stored at -80°C prior to DNA extraction. High-molecular weight DNA was isolated from leaves following a CTAB protocol (Callahan *et al.*, 2021) with slight modifications. Extraction and organelle wash buffers were supplemented with proteinase K (Ambion) – 3 ug per 5 ml and 8 ul per 4 ml respectively. Treatment of DNA with an RNAse cocktail (Ambion) was conducted in a TE buffer containing 1 M NaCl. The quality and integrity of the DNA were evaluated using a Qubit 2 Fluorometer (Thermo Fisher Scientific Inc., USA) and a NanoDrop ND-8000 (Thermo Fisher Scientific Inc., USA), followed by pulse-field electrophoresis (Sage Science, Inc). DNA underwent library preparation, followed by two rounds of PacBio Sequel II HiFi circular consensus sequencing (Novogene), yielding 34.45 Gb of sequence with an average read length of 20,753 bases. Reads with over 90% of length matching the publicly available *Q. dentata* chloroplast genome sequence NCBI accession NC_039725.1 (Hu *et al.*, 2019) were removed. DNA from flash frozen leaves was also used for Hi-C library preparation (Phase Genomics), which was sequenced on an Illumina NovaSeq S2 lane at the North Carolina State University Genomic Sciences Laboratory and yielded 402.8 million read pairs. The quality of the Hi-C data was assessed with hic_qc (Phase Genomics, 2021).

**Reference genome assembly and annotation**

The cleaned PacBio and Hi-C Illumina reads were assembled with Hifiasm v.0.16.1 (-s = .50) (Cheng *et al.*, 2021), yielding a primary (hapA) and an alternate (hapB) assembly. The assemblies were compared the *Quercus dentata* chloroplast (NCBI RefSeq NC_039725.1) and *Quercus acutissima* mitochondria (NCBI Genbank MZ636519.1) with minimap2-2.24 (Li, 2018)). A total of eight contigs with greater than 90% length mapped to the chloroplast or the mitochondrial genomes were removed. The plastome and mitochondrial genome **(See below)** of the MM1 individual were assembled and annotated separately. Hi-C data was mapped to the assembled contigs with Juicer 1.6 (Durand *et al.*, 2016) then ordered and oriented into chromosomes with 3d-dna v201008 (Dudchenko *et al.*, 2017) and the parameters "--editor-repeat-coverage 8", " --editor-coarse-resolution 50000", and "--editor-coarse-region 250000". Chromosomes were reviewed in Juicebox v1.11.08 (Robinson *et al.*, 2018) and five putative inversions between the two haplotypes with low Hi-C support for one haplotype were manually corrected. To assess genome quality, BUSCO v5.2.2 was run on the primary and alternate haplotype assemblies using the embryophyta_odb10 database, which contains 1,614 BUSCOs (Manni *et al.*, 2021).

We constructed a genetic map based on 184 full- and half-sibling *Q. alba* individuals (**See below**) and compared 177 genetic map markers with available sequences to the chromosomes and unplaced scaffold sequences with BLAST+ (Camacho *et al.*, 2009). Markers with multiple equivalent e-value hits at different locations, which likely represent markers derived from multi-member gene families, were removed from the analysis. The hapA and hapB chromosomes were numbered and oriented following the genetic map, which also follows the *Q. robur* and *Q. lobata* genome naming and orientation convention. Visualization of the alignment of the genetic map and the chromosome assemblies was produced by RIdeogram (Hao *et al.*, 2020).

Repetitive DNA and gene annotation were completed independently for the hapA and hapB assemblies. The full assemblies were analyzed by RepeatModeler v2.0.3 (Flynn *et al.*, 2020). To ensure that the *de novo* repeats do not overlap with protein coding genes, the repeats were removed if they had BLAST matches with a minimum e-value of 1e-30 to any genes in the *Quercus rubra* v2.1 genome assembly available at Phytozome (Kapoor *et al.*, 2023). This removed about 10% of repeats called by RepeatModeler. To call LTRs, a combination of LTRHarvest (Ellinghaus *et al.*, 2008) and LTR_Finder (Xu & Wang, 2007) results were provided to LTR retriever 2.9.0 (Ou & Jiang, 2018), following the recommended procedures of LTR retriever and using default parameters. LTR retriever outputs the LAI (LTR Assembly Index). The LTR repeats were also screened for matches to *Quercus rubra* protein coding genes as described above. Soft masking of the assemblies was performed with RepeatMasker (Smit *et al.*, 2004) using the filtered RepeatModeler output, the filtered LTR retriever output, and sequences from the Dfam database of repeats from eudicotyledons only (Hubley *et al.*, 2016).

The synteny and structural variation of the primary and alternate haplotypes were analyzed by the Synteny and Rearrangement Identifier (SyRI) v1.5.4 (Goel *et al.*, 2019) based on a mapping of the alternate assembly to the primary assembly with minimap2 v2.24 (Li, 2018). Visualizations of the SyRI results were constructed by plotsr v0.5.1 (Goel & Schneeberger, 2022).

RNA sequencing was performed on leaves from MM1; leaves, buds, bark, roots, and wood samples from a seedling originally from Oldham Co., KY and grown at the Kentucky Division of Forestry Nursery; an ungerminated acorn from Ashley County, Arkansas; and a germinating acorn from Knox County, Tennessee. RNA was extracted following a previously published protocol for oak (Soltani *et al.*, 2020), and sequenced by Novogene on an Illumina instrument. A set of four accessions of RNA sequenced by the 454 platform were downloaded from NCBI (SRR006309 - SRR006312). Illumina and 454 sequenced RNA data were mapped to each haplotype assembly with STAR 2.7.9a (Dobin *et al.*, 2013). BRAKER 2.1.6 (Bruna *et al.*, 2021) was run with the full set of RNASeq data. A second BRAKER annotation round was completed using predicted proteins from *Q. rubra* v2.1 and *Q. robur* version PM1N mapped with ProtHint 2.6.0 (Bruna *et al.*, 2020). TSEBRA was used to combine the two BRAKER annotation sets (Gabriel *et al.*, 2021). This set of combined annotations was filtered by gFACs v1.1.2 with parameters "--unique-genes-only" and "--rem-genes-without-start-and-stop-codon" (Caballero & Wegrzyn, 2019). The genes were functionally annotated with EnTAP v0.10.8 (Hart *et al.*, 2020), which was configured to compare sequences to Uniprot's SwissProt and TrEMBL protein databases (Bairoch *et al.*, 2005), with the latter filtered to only plant proteins. Genes that were on an unplaced scaffold and lacked annotation by EnTAP were removed from the gene set. HTSeq was used to assess the number of mapped RNASeq reads overlapping a gene annotation (Anders *et al.*, 2015). Bedtools v2.31.0 (Quinlan & Hall, 2010) was used to assess the number of gene bodies and gene exonic regions overlapped a structural variant identified between hapA and hapB.

The locations of rDNA arrays in hapA and hapB were identified with rnammer (Lagesen *et al.*, 2007). Fluorescence *in situ* hybridization (FISH) with rDNA oligonucleotide probes was conducted following the methods of Kapoor et al. (2023). R-gene domains and categories were determined using DRAGO2 (Calle García *et al.*, 2022). Gene duplication mechanisms were determined by the MCScanX duplicate gene classifier tool (Wang *et al.*, 2012) of the white oak genome hapA against itself, resulting in the assignment of five categories of gene expansion type: (1) whole genome /segmental duplication where a set of collinear genes are found duplicated in collinear blocks, (2) tandem duplication where duplicate genes are adjacent to each other, (3) proximal duplication where genes are not adjacent but are in nearby chromosomal regions, (4) dispersed duplications where duplicated genes are found but do not fit the criteria for other categories, and (5) singleton genes that have no identified duplication from other genes.

**Assembly and annotation of the *Querucs alba* MM1 plastome**

We generated a draft assembly of the plastid genome from the MM1 individual with HiC library Illumina reads trimmed with Skewer v0.2.2 **(Jiang et al 2014)** and GetOrganelle v1.7.7.0 **(Jin et al., 2020)** with the setting "-R 10". This resulted in a circular plastome assembly, which was 161,104 base pairs in length, with four possible orientations, one of which matched the conventional angiosperm plastome structure and was selected for further curation. In order to correct any assembly errors introduced by the use of HiC library reads, we mapped PacBio CCS reads to this draft assembly. First, we used MiniMap2 (Li, 2018) to map all PacBio reads from the MM1 individual **(See above)** to a *Querucs alba* plastome assembly (sample QalbaRNA_S24) generated with GetOrganelle in the same way, but with standard Illumina library reads. Reads that mapped to the QalbaRNA_S24 plastome assembly were then mapped to the draft MM1 assembly using MiniMap2 within Geneious Prime 2023, which resulted in 26,203 reads mapping to the draft MM1 assembly. We extracted the consensus sequence based on the mapped reads, which resulted in a sequence that was 161,180 base pairs in length and included two "N" calls at sites 35100 and 66311. These "N"s preceded one multi-A run and one multi-T run, respectively, and appeared to be the result of differing consensus lengths of these runs among PacBio CCS reads. These two "N" sites were removed from the draft plastome consensus sequence. Therefore, the final plastome sequence was 161,178 base pairs in length. The contents of the final sequences were: A: 50,222 (31.2%); T: 51,583 (32.0%); C: 30,234 (18.8%); G: 29,138 (18.1%). There was one site with several mapped CCS reads supporting both G and C calls, for which the IUPAC code "K" appears in the final assembly. We used the GeSeq program **(Tillich et al., 2017)** from the chlorobox web server to annotate the MM1 plastome assembly. Annotations were manually removed in cases where an annotation for a gene fragment fell within another gene. The final annotated plastome was visualized with OGDraw (Greiner *et al.*, 2019).

#### Identification and annotation of *Quercus alba* (MM1) mitochondrial genome scaffolds

Two scaffolds generated with HiFasm, which were 321,000 and 95,221 base pairs in length, respectively, were identified as containing mitochondrial genome sequences based on alignment with minimap2 (**See above**). The contents of these scaffolds was A: 111,990 (27.1%); G: 95,547 (23.1%); C: 94,053 (22.8%); T: 111,631 (27.0%); N: 3,000 (0.7%). These scaffolds were annotated with the live annotate and predict feature in Geneious Prime 2023 based on publicly available annotated mitochondrial genome sequences from *Q. acutissima* (Genbank accession MZ636519.1) and *Q. variabilis* (Genbank accession MN199236.1), with a 70% similarity threshold and the best match option. For visualization, the two scaffolds were concatenated with 100 "?"s inserted as spacers at junctions between scaffolds and input into OGDraw (Greiner *et al.*, 2019).

#### Genetic map construction

Progeny derived from the open-pollinated mother trees WO1 (~700 acorns) were maintained at the Ames Plantations (Fayette County, Tennessee). A preliminary parental test with five SSRs identified 14% full-sibs among WO1-derived seedlings. Of these, 184 progeny from the WO1 mother tree were selected for further genotyping and map construction. Genotyping was conducted using the Sequenom genotyping massarray method (Bradic et al., 2011). Markers segregating in the 1:1 and 1:2:1 ratios were encoded as "*lm x ll*" and "*hk x hk*", respectively. The female parent map was constructed under the cross-pollinated population type with JoinMap4.1 (Van Ooijen, 2006). Markers were assigned to 12 linkage groups at logarithm of odds (LOD) score > 4.0. Within linkage groups the marker order was calculated with the regression mapping algorithm and Kosambi function with the following parameters: maximum recombination frequency 0.25, minimum LOD of 1.0 and goodness-of-fit jump threshold for removing loci of 5.0. For map saturation, we used "fixed map function" and added 67 SNPs genotyped in 30 individuals in an independent white oak mapping project (Di Wu, personal communication). Linkage groups were named and oriented against the *Q. robur* genetic map by Bodenes *et al.* (2016) and the *Q. rubra* map by Konar *et al* (2017). Graphical representation of the map was drawn with MapChart 3.0 (Voorips, 2002).

#### Population genomics sampling and DNA Sequencing

We sampled 16 *Q. alba* individuals (four individuals from Wisconsin, Ohio, Indiana, and Mississippi), growing as part of a provenance trial near Vallonia, Indiana (**Table S1**). For each, we extracted DNA with the DNAeasy Plant Pro kit (Qiagen) following the manufacturer's protocol, except that we ground 100mg of fresh leaves with liquid nitrogen and mixed the powder with 450 ul solution CD1 and 50 ul solution PS in a 2 ml tissue disruption tube, which was then vortexed at max speed for 5 minutes. We also conducted DNA sequencing on two *Q. alba* individuals from Kentucky: an individual that was used for RNA-seq to annotate the *Q. alba* genome (seedling of Oldham County, Kentucky origin) and the "Presidents Tree" from the University of Kentucky campus in Lexington, KY (**Table S1**). Illumina (San Diego, CA) 150-bp paired-end reads were generated for all 18 samples. We also included the MM1 tree in our population genetics sampling – sequence data for which this individual were two runs of PacBio HiFi long reads described above, which were combined prior to analysis.

**Population genomics read processing and genotyping**

All Illumina reads were trimmed for adapters and quality with Skewer v0.2.2 (Jiang *et al.*, 2014) and quality checked with Fastqc (Andrews, 2010). HiFi long reads that contained adapters were removed with HiFiAdapterFilt (Sim *et al.*, 2022) and reads that had fewer than four sequencing passes were removed with a custom python script. Next, all reads were aligned to hapA of the *Q. alba* reference genome with BWA-MEM v0.7.12 (Li & Durbin, 2009). The resulting bam files were sorted with Samtools v1.15.1 (Li *et al.*, 2009). Read groups were added to each bam with Picard v2.27.3 (Broad Institute, 2023) and were indexed with samtools. Next, each bam was divided into 13 separate files (i.e. one for each of the 12 chromosomes and one for unplaced scaffolds) per sample with the *view* command in samtools to facilitate downstream genotyping. All mapped short reads were then processed with the *MarkDuplicates* program in Picard v2.21.9 as implemented in GATK v4.1.7 (McKenna *et al.*, 2010; DePristo *et al.*, 2011; Van Der Auwera *et al.*, 2013). For HiFi reads, duplicates were marked with Picard v2.27.3. HaplotypeCaller was then run on all mapped reads to produce separate gvcfs files for each chromosome for each sample, which were combined per chromosome and used for variant calling with the *GenotypeGVCF* command in GATK. Variant calls for 19 individuals and all 12 chromosomes were then combined into a single vcf file with the *GatherVCFs* command in GATK. Unplaced scaffolds were not included in further analyses and made up only 3.14% of the genome assembly. Variant calls were soft filtered with the settings QD < 2.0, FS > 60.0, MQ < 40.0, SOR > 4.0, MQRankSum < -12.5, and ReadPosRankSum < -8.0 with the *VariantFiltration* command in GATK and then removed, along with indels, from downstream analyses with the *SelectVariants* command in GATK and the options "--select-type-to-include SNP" and "--exclude-filtered true" to produce a "filtered SNP dataset" used in several downstream analyses.

**Population genomics clustering analyses**

To produce a dataset for genetic clustering, the filtered SNP dataset was further processed using PLINK v2.0 and the options "--min-alleles 2", "--max-alleles 2", "--snps-only just-acgt", "--geno 0", and "--thin-count 10000" to produce a dataset of 10,000 randomly selected, bialleleic SNPs with no missing genotype calls for any of the 19 *Q. alba* samples (Purcell *et al.*, 2007). These SNP data were reformatted with PLINK v1.9 and the option "--recode structure" and used as the input for analysis in *Structure* v2.3.4 (Pritchard *et al.*, 2000). We ran 10 replicate *Structure* runs each for each value of K from 1 to 10 with a unique random seed for each replicate. The results were summarized with the CLUMPAK online server and default settings (Kopelman *et al.*, 2015). We considered K=3 as our focal value of K in our results and discussion, since CLUMPAK suggested high similarity (91.2%) among all replicates at this value, but not for higher values of K and results for K=3 were qualitatively similar to results from other analyses (see Results).

**Population genomics principal component analyses and population genetic statistics**

We generated a biallelic SNP dataset for principal component analysis (PCA) and estimating population genetic statistics and by processing the filtered SNP dataset with PLINK v2.0 and the options "--snps-only", "--min-alleles 2", and "--max-alleles 2" (Purcell *et al.*, 2007). In order to prepare this dataset for PCA analysis, we assigned unique variant IDs to all SNPs with a custom python script and calculated allele frequencies with the "--freq" command in PLINK v2.0. We then conducted a PCA with PLINK v2.0 and the options "--read-freqs" and "--pca". We visualized the first two principal components, which correspond to the first two eigenvectors, using the ggplot2 library and a custom R script (Wickham, 2016). The percent variance explained by each principal component was calculated in R using the eigenvalues output by PLINK v2.0.

We calculated pairwise $F_{ST}$ values among "populations" by grouping individuals in three ways. First, we grouped samples based on their state of origin and refer to these as the "by state analyses". Second, we considered all individuals from Indiana, Ohio, and Kentucky as a single population, since these individuals originated geographically close to one another and clustered together in the PCA analysis, and compared this to groups of samples from Wisconsin and Mississippi. We refer to these as the "geographically clustered analyses". Third we grouped samples based on their major cluster assignment inferred in *Structure* when K=3, which we refer to as the "genetically clustered analyses". We calculated $F_{ST}$ using both the (Hudson *et al.*, 1992) and (Weir & Cockerham, 1984) estimators on a per-SNP basis with PLINK v2.0 and then calculated genome wide averages with a custom Python script. With the same biallelic SNP dataset and group delineations, we calculated nucleotide diversity ($\pi$) at all SNPs using VCFtools v0.1.13 (Danecek *et al.*, 2011). We then calculated genome-wide averages of $\pi$ with a custom Python script that accounted for sites that did not vary within the group, sites with 3+ alleles, and sites that had only missing data within the group (but that were variable when all 19 samples were considered together). Any sites that were not characterized as SNPs in the filtered SNP dataset with all 19 individuals (e.g. sites with only missing data, potential variants that did not pass filtering) were assumed to be invariable for the purpose of these calculations.

**Phylogenomics sequencing, read acquisition, and genotyping**

We assembled a phylogenomic dataset with data from several sources including new sequencing and the NCBI SRA (**Table S1**). We conducted whole genome resequencing (as described above) for samples from *Quercus* individuals growing in The Arboretum at the University of Kentucky (State Botanical Garden of Kentucky), Lexington, KY representing seven species in the white oak clade (section *Quercus*), the provenances of which were various locations within the state of Kentucky, USA. We obtained/generated whole genome resequencing data for two red oaks (*Q. rubra*) associated with the red oak genome project (Kapoor *et al.*, 2023). In addition, we obtained whole genome resequencing data with full base quality scores for 15 samples from the NCBI Sequence Read Archive (SRA) using the fasterq-dump command from the sra-toolkit v3.0.0 (NCBI, 2023). The SRA samples comprised 14 species, including an additional high-coverage sequencing run for an individual of *Q. rubra* as well as an individual of *Lithocarpus longipedicellatus* (**Table S1**). The 19 samples of *Q. alba* described above were also included in the phylogenomic dataset construction. All Illumina reads were trimmed with Skewer v0.2.2 (Jiang *et al.*, 2014). The phylogenomic dataset included 43 individuals in total.

We employed a pseudo-reference approach to produce genome sequences for each individual with several iterations of site masking. Variant calling was conducted for a single chromosome of a single individual at a time, rather than jointly genotyping any individuals. First, sorted BAM files with duplicate reads marked with *MarkDuplicates* (see above) were used as input for the *mpileup* command in bcftools (Li, 2011) and the options "-Ou", "-a FORMAT/AD,FORMAT/DP", "--skip-indels", "--min-BQ 15" and "--min-MQ 15" followed by the *call* command in bcftools with the options "-m", "--gvcf 0" and "-Oz". These commands produced a genomic vcf file for each sample that included all invariable sites, even those with no mapping reads. Next, vcftools was used to remove sites from gvcfs which had coverage that was greater than two times the average coverage for that sample (calculated beforehand with the *depth* command is Samtools) as well as sites without a genotype call for that sample with the options "--maxDP" and "--geno 1" (Danecek *et al.*, 2011). The resulting file was indexed with BCFtools and used as input for the *consensus* command in BCFtools with the options "--haplotype I", "--absent '?' ", "--missing 'n' ", and "--mark-del '-' ". This resulted in a fasta file for each chromosome for each sample, with heterozygous genotypes coded with IUPAC ambiguity codes. Each chromosome sequence was then processed with a custom Python script that randomly selected one of the two alleles at heterozygous sites, in order to generate haploid sequences, and all sites identified as repetitive DNA in hapA of the reference genome by RepeatMasker (Smit *et al.*, 2004) hard masked as "N"s. For each individual, the 12 chromosome sequences were then combined into one file, resulting in a single fasta file per sample.

As a final step to prepare our pseudo-reference sequences, we used the Referee package (Thomas & Hahn, 2019) to calculate genotype quality scores for each site and masked sites where the base call was not supported. For diploid organisms, there are 10 possible genotypes for any given site. Referee compares the likelihoods of genotypes that include the (pseudo-)reference base against the possible genotypes that do not include that base. To prepare input files for Referee, we indexed files as necessary, mapped the corresponding trimmed reads to each pseudo-reference, sorted the resulting bam files, and marked duplicate reads with Picard v2.27.3 as above. We then generated an mpileup file with bcftools which we used as input for Referee with the options "--pileup" and "--mapq". Finally, we masked sites in each pseudo-reference where the likelihood was better for genotypes that did not include the called base (i.e. sites where the called base was not supported).

The final phylogenomic matrix consisted of 327,242,758 aligned sites that were not masked in all samples. Among these sites, 1,564,543,424 nucleotides (12.6%) were masked in individuals because of missing genotype calls, excessively high coverage, or lack of support from the Referee pipeline. There were 123,757,882 sites (37.8%) with genotypes for all samples and a further 44,019,896 sites (13.5%) with genotypes for all but one individual. There were 323,540,424 sites (98.9%) with genotypes for at least four individuals and 24,366,683 sites (7.4%) that were parsimony informative. There were a total of 770,665,549 aligned sites in the dataset (including sites that were masked in all samples such as repetitive DNA regions).

We constructed 5 kb non-overlapping windows, with the same taxa as the phylogenetic matrix described above except that we included only two *Q. alba* individuals – one sample from Wisconsin and one from Mississippi with the highest effective coverage (i.e. PS96_S14 and PS102_S5). Using the same masked sequences used to construct the whole genome matrix, we generated 5kb window alignments, excluding windows where >50% of sites were masked in all included samples. This resulted in 64,761 windows. All "?" symbols were changed to "N" in all window alignments. Next, to retain only windows with a high proportion of non-missing data, we assessed data occupancy by using the pxclsq command in phyx **[Brown et al., 2017]** and kept only windows for which ≥80% of sites contained non-missing data in ≥80% of taxa, which resulted in 12,091 windows that met these criteria. To remove individual sequences with a high proportion of missing data, sequences with fewer than 1000 bp of non-missing/ambiguous data (i.e. 20% data occupancy) were removed from window alignments. This resulted in a final window dataset with 11,930 (98.7%) windows that included all 21 taxa, 146 (1.21%) windows with 20 taxa, six (0.05%) windows with 19 taxa, eight (0.07%) windows with 18 taxa, and one window with 17 taxa.

#### Phylogenomic analysis

To infer phylogenetic relationships, we selected a subset of the pseudo-reference genomes described above, prioritizing samples with at least 10X sequence coverage genome-wide (**Table S1**). We chose a single individual per species, except for *Q. alba*, for which we included all individuals and generated a matrix of whole genome alignments of all 12 chromosomes, excluding unplaced scaffolds. The matrix included 37 individuals from 19 species of *Quercus* plus one individual of *Lithocarpus* as an outgroup. Because all pseudo-reference sequences were generated relative to hapA of the *Q. alba* reference genome, there was no need to align sequences prior to phylogenetic analysis. We estimated a maximum likelihood phylogenetic tree with IQ-TREE v2.2.0 with a single GTR+G model of evolution; site concordance factors (sCF) were calculated with the "--scf" option and 1000 quartet replicates (Minh *et al.*, 2020b,a). For visualization and divergence time estimation, a version of the tree with *Q. alba* represented by only MM1 was generated using the *pxrmt* command in phyx (Brown *et al.*, 2017). Window trees for the 12091 genomic windows described above were estimated with IQ-Tree v2.0.7 and the GTR+G model and 1000 ultra-fast bootstraps **(Minh et al., 2020; Hoang et al., 2018]**. We estimated a species tree from all resulting window trees with ASTRAL v5.7.7 **(Zhang et al., 2018**) and default settings after trimming PS102_S5 from window trees with phyx, so that *Q. alba* was represented by a single individual. We assess gene concordance factors (gCF) values for the maximum likelihood and ASTRAL trees with IQ-TREE v2.2.0, the "-gcf" option, the 12,091 window trees, and GNU parallel **(Minh et al 2020b; Tange 2018)** after trimming all trees so that *Q. alba* was represented by only sample PS96_S14. We used the ape package (Paradis & Schliep, 2019) in R (R Core Team, 2013) to calculate the Robinson-Foulds distances (Robinson & Foulds, 1981) for the 11,930 trimmed window trees with full taxon sampling compared to the trimmed maximum likelihood and ASTRAL trees.

In order to investigate genetic variation that is shared between *Quercus alba* and other oak species, we further analyzed the masked pseudo-reference alignment described above. First, we determined which sites were variable within the 19 individuals of *Q. alba*, with and without excluding singletons (i.e. not counting a site as variable if only one *Q. alba* individual differed from the others). Then, for each site that was variable in *Q. alba*, we investigated whether that site was also variable among other North American white oak species (which were inferred to form a clade with *Q. alba* in our analyses; see below), excluding *Q. alba*, and whether that site was variable among all white oak species in our sampling. Variable sites within *Q. alba* for which at least two of the same alleles were also variable among other white oak species, were counted as a shared variable site. Because we considered haploid sequences for this analysis, our results are conservative estimates of the amount of genetic variation shared among species.

We generated an ultrametric phylogeny with branch lengths scaled to time using the penalized likelihood approach as implemented in treePL (Sanderson, 2002; Smith & O'Meara, 2012). Prior to dating the phylogenetic tree, we trimmed the tree to remove the outgroup and all *Q. alba* except MM1 using the *pxrmt* command in phyx (Brown *et al.*, 2017). To account for ancestral polymorphism and to transform our tree from one that represented genetic divergence among individuals to one that represented divergences among species (Edwards & Beerli, 2000), we used our estimate of nucleotide diversity ($\pi$) calculated for *Q. alba* (*i.e.*, 0.012) and subtracted $\pi/2$ from each terminal branch. Then, we dated the resulting tree using a calibration for the crown age of *Quercus* at 56 Ma (Hofmann, 2010; Hofmann *et al.*, 2011; Hipp *et al.*, 2020) with priming and optimization with the *randomcv* option in treePL. Cross validation resulted in an optimal smoothing parameter of $1x10^{-4}$. For comparison, we also used the same methods to date the maximum likelihood tree without corrections to branch lengths, which resulted in an optimal smoothing parameter of $1x10^{9}$. The full configuration file with all parameters is available from the Dryad submission accompanying this article.

#### Plastid and mitochondrial dataset generation and analysis.

We used Illumina reads trimmed with Skewer v0.2.2 (Jiang *et al*., 2014) and the program GetOrganelle (Jin *et al*., 2020) to assemble plastid and mitochondrial genomes for 42 *Quercus* individuals and one *Lithocarpus* under a variety of settings (**Table S1**). Complete plastomes assemblies typically have two possible configurations with alternate orientations of the short single copy (SSC) region. For each assembly, we annotated and extracted gene sequences from the first configuration output by GetOrganelle with PhyloHerb (Cai *et al.*, 2022) with the "-m ortho"' option and the "-mito" option for mitochondrial assemblies. We were not able to generate a mitochondrial assembly for three samples **(Table S1)**, therefore, downstream analyses for the mitochondrial genome dataset included 40 individuals. Gene sequences were individually aligned with the default algorithm in MAFFT v7.505 (Katoh *et al.*, 2002; Katoh & Standley, 2013) and the options "--maxiterate 1000" and "--adjustdirection". Alignments were then individually examined with Aliview (Larsson, 2014) and poorly aligned regions (that likely represented annotation errors or short structural differences) were manually masked with 'N's. Some mitochondrial genes were excluded from further downstream analysis due to pervasive alignment issues. We concatenated the 87 curated plastid gene alignments totalling 93,946 sites (6.2% missing data, 610 parsimony-informative sites) with the *pxcat* command in phyx (Brown *et al.*, 2017) and estimated a tree with IQ-TREE v2.2.0 (Nguyen *et al.*, 2014; Mihn *et al.*, 2020) under a single GTR+G model. Similarly, we concatenated the 36 curated mitochondrial gene alignments totalling 33,257 sites (13.1% missing data, 112 parsimony-informative sites) and estimated a tree with IQ-TREE under a single GTR+G model. Trees were visualized with Figtree (http://tree.bio.ed.ac.uk/software/figtree/) and the *cophylo* command from Phytools (Revell, 2012) using a custom R script. Individual curated and uncurated gene alignments as well as concatenated alignments are available from the repository submission that accompanies this article.

#### Comparative Genomics

Syntenic structure for ten genomes was assessed using SyRI as described above, with *Q. alb*a hapA as the reference. The genomes were: *Q. lobata* version 3.0 (Sork *et al.*, 2022), *Q. robur* version dhQueRobu3.1 (The Darwin Tree of Life Project Consortium *et al.*, 2022), *Q. mongolica* version GCA_011696235 (Ai *et al.*, 2022), *Q. rubra* version version 2.1 (Kapoor *et al.*, 2023), *Q. gilva* version GCA_023621385.1 (Zhou *et al.*, 2022), *Q. acutissima* version GWHBGBO00000000 (Fu *et al.*, 2022), *Q. variabilis* version CNP0003390 (Han *et al.*, 2022), *Q. aquifolioides* version GCA_019022515 (publicly available from NCBI), *Castanea mollissima* version GCA_014183005.1 (Wang *et al.*, 2020), *Castanopsis tibetana* version CNSA_CNP0001714 (Sun *et al.*, 2022).

A syntenic block analysis was conducted by using OrthoFinder v2.5.4 (Emms & Kelly, 2019) on primary proteins, followed by syntenic block identification and duplicate gene classification by MCScanX (Wang *et al.*, 2012). The analysis included eleven species from the order Fagales in addition to *Quercus alba*: (Fagaceae) *Castanea mollissima* (Wang *et al.*, 2020), *Castanopsis tibetana* (Sun *et al.*, 2022), *Fagus sylvatica* (Mishra *et al.*, 2021), *Quercus lobata* (Sork *et al.*, 2022), *Quercus mongolica* (Ai *et al.*, 2022), and *Quercus robur* dhQueRobu3.1; (Betulaceae) *Carpinus fangiana* (Yang *et al.*, 2020), *Corylus avellana* (Lucas *et al.*, 2021); (Juglandaceae) *Carya illinoinensis* (Lovell *et al.*, 2021), *Juglans regia* (Martínez-García *et al.*, 2016); and (Myricaceae) *Morella rubra* (Jia *et al.*, 2019). *Prunus persica*, a species from the order Rosales, was also included (International Peach Genome Initiative *et al.*, 2013).

Gene families were determined in hapA and primary protein sets from seven of the *Quercus* species described above by running GeneSpace (Lovell *et al.*, 2022), which uses OrthoFinder v2.5.4 and MCScanX. Gene duplication mechanisms were determined by the MCScanX duplicate gene classifier tool (Wang *et al.*, 2012) with the same methods as for the *Q. alba* reference genome. Expansion and contraction of gene families was determined by CAFE5 (Mendes *et al.*, 2020) with the base model.

**Supporting Notes**

**Notes S1.** Details of differences in Tc1/mariner family of repeats in HapA and HapB

The DNA transposon Tc1/mariner superfamily (labeled "Tc1-IS630-Pogo" by RepeatMasker) constituted 0.08% of hapA but 1.43% of hapB (**Table S6**). This family of repeats consisted of three identified repeat types: Stowaway, Tc1, and Mariner. Stowaway repeats, MITES (miniature inverted repeat transposable elements) found in many plant genomes (Feng, 2003; González & Petrov, 2009), are similar between the two white oak haplotypes. HapA has 434 kb and hapB has 423 kb of Stowaway repeats, and in both cases the repeats are dispersed across chromosomes with less than 3 kb annotated in unplaced scaffolds. In contrast, Tc1 is only found in hapA while Mariner is only found in hapB. Tc1 repeats span 202 kb of HapA, dispersed across all 12 chromosomes (<2 kb on unplaced scaffolds). In contrast, hapB has 10.9 Mb of Mariner repeats, which is 1.38% of the total hapB content. A total of 4.9 Mb of Mariner repeats (45%) are found in hapB unplaced scaffolds and make up 20.6% of total unplaced base pairs. These results suggest that this repeat family was not able to be accurately scaffolded and should be further explored in future genome versions.

**Notes S2.** Additional discussion of phylogenetic relationships.

Hipp *et al*. (2020) recovered *Quercus alba* + *Q. montana* + *Q. michauxii*, which they termed the "Albae" clade. However, we recovered *Q. alba* and *Q. montana* as sister taxa, with *Q. michauxii* more closely related to *Q. bicolor* and *Q. lyrata* - members of the "Prinoids" clade *sensu* Hipp *et al*. (2020). That study also found the Albae clade to be sister to the "Roburoids" (represented by *Q. mongolica*, *Q. robur*, and *Q. petraea* in our sampling), whereas we found all North American white oaks (including *Q. lobata* and *Q. stellata*) to form a clade that was sister to the Roburoids. Within subgenus *Cerris*, we found *Q. chenii* and *Q. acutissima* to be sister in our sampling, with *Q. variabilis* sister to those, whereas Hipp *et al.* (2020) found *Q. chenii* to be sister to *Q. variabilis*.

**Supporting Figures**

**Figure S1.** Summary of taxa included in each comparative genomics analysis. Filled circles indicate taxa were included in the analysis, empty circles indicate the taxon was not included. Family names are included for taxa outside Fagaceae.

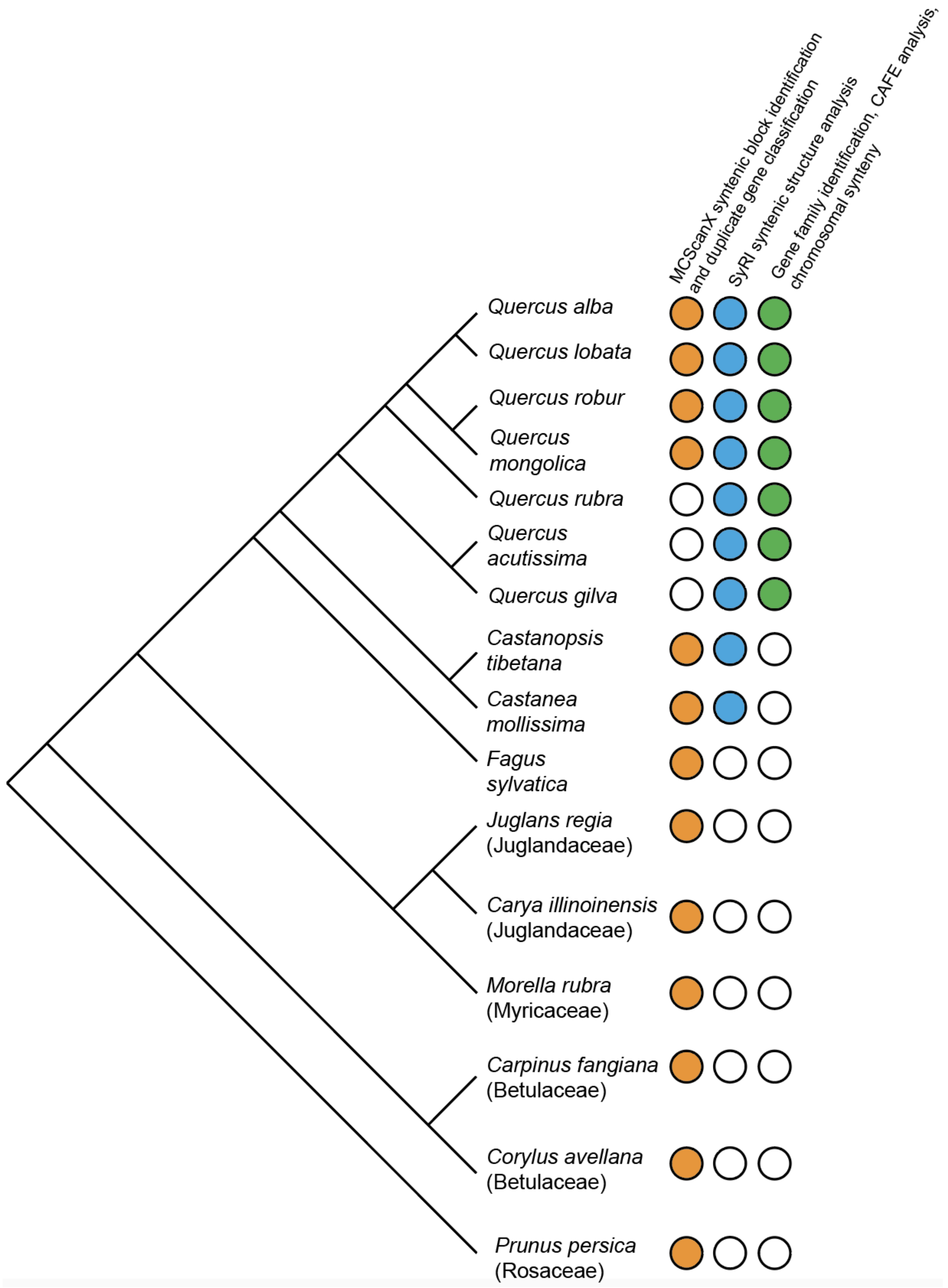

**Figure S2.** The *Quercus alba* (MM1) plastome. The plastome was assembled from Illumina HiC library reads with GetOrganelle, followed by mapping of PacBio CCS reads with minimap2 and consensus sequence calling.

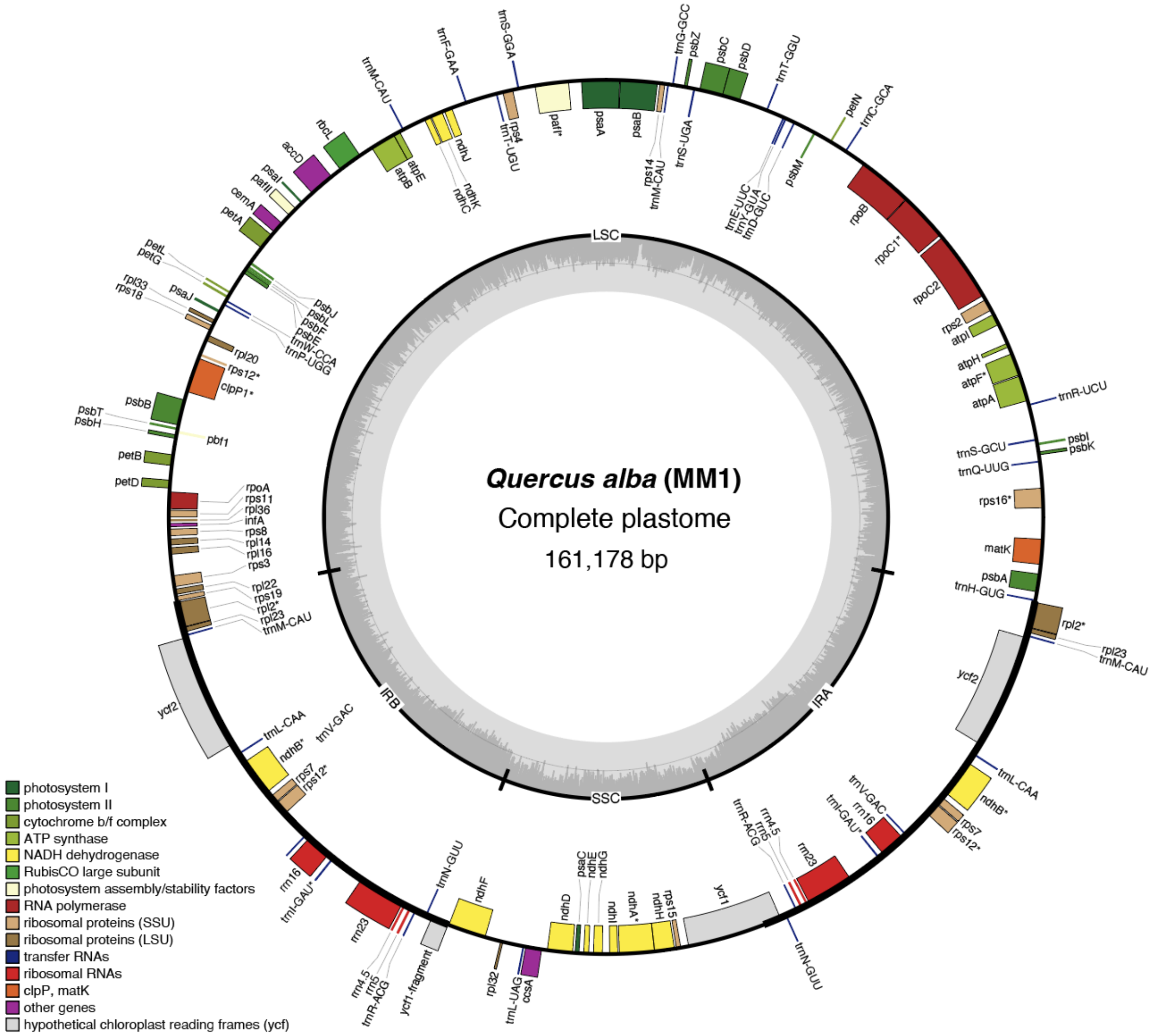

**Figure S3.** Annotated, circularized draft assembly of the *Quercus alba* (MM1) mitochondrial genome, based on two scaffolds recovered with HiPhasm (see main text and Methods S1). Scaffolds are presented here in arbitrary orientations. Approximate breakpoints between scaffolds are denoted with dashed lines. **The draft genome assembly includes 413,221 base pairs of nucleotides, not including 'N's or '?'s.

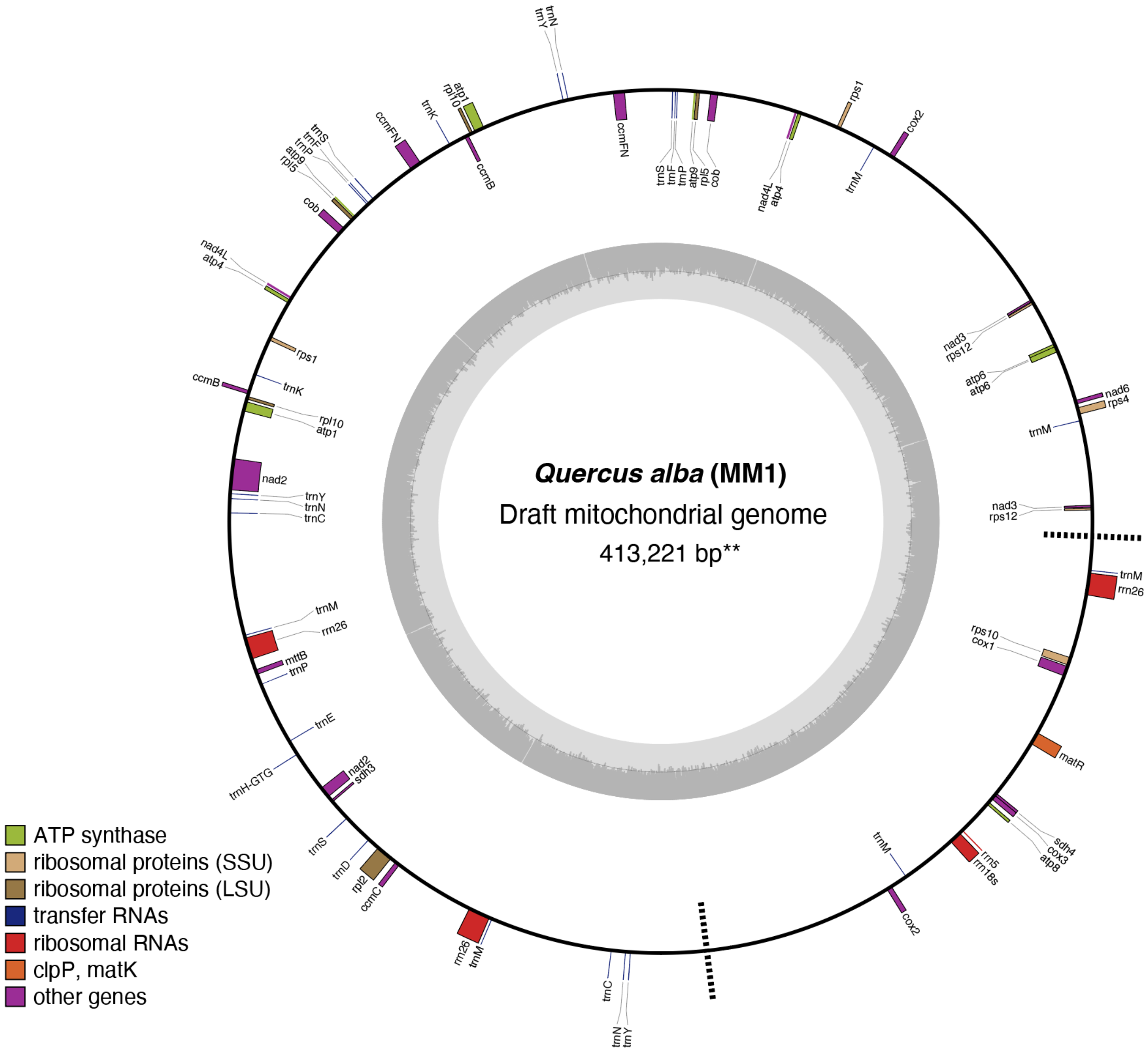

**Figure S4.** Female WO1 map composed of 181 SNP markers.

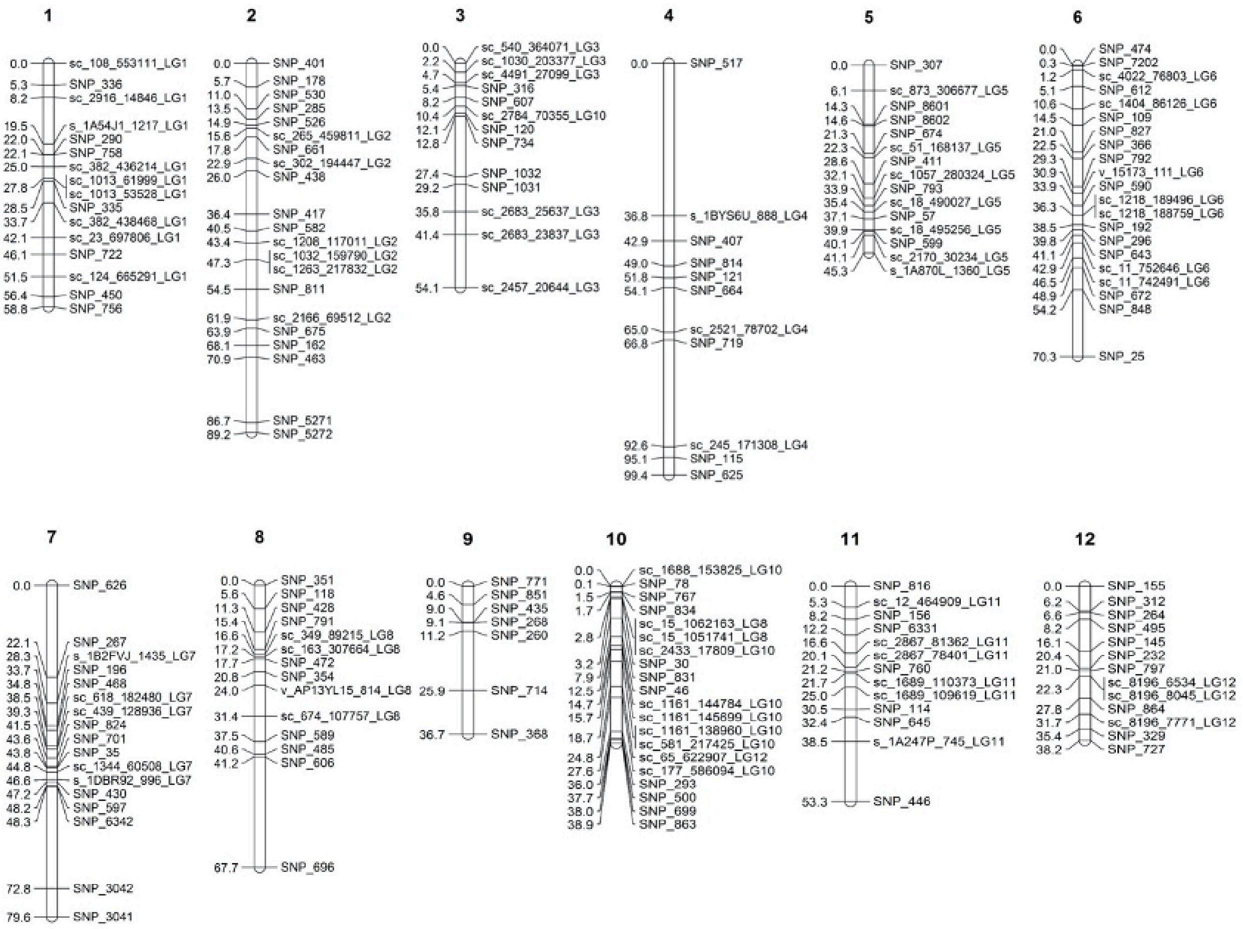

**Figure S5**. Fluorescence in situ hybridization (FISH) of white oak chromosome spreads reveals two pairs of 35S (green) and one pair of 5S (red) rDNA signals. A prometaphase spread and an interphase nucleus. Each of the major 35S rDNA bearing chromosomes showed a pair of signals that indicates the major site has two blocks of signals. Enlarged images of the major rDNA bearing chromosomes shown in inserts. The arrowheads in inserts point at the respective satellite. A metaphase spread with 35S and 5S rDNA signals in displayed in the main manuscript (Figure 2b)

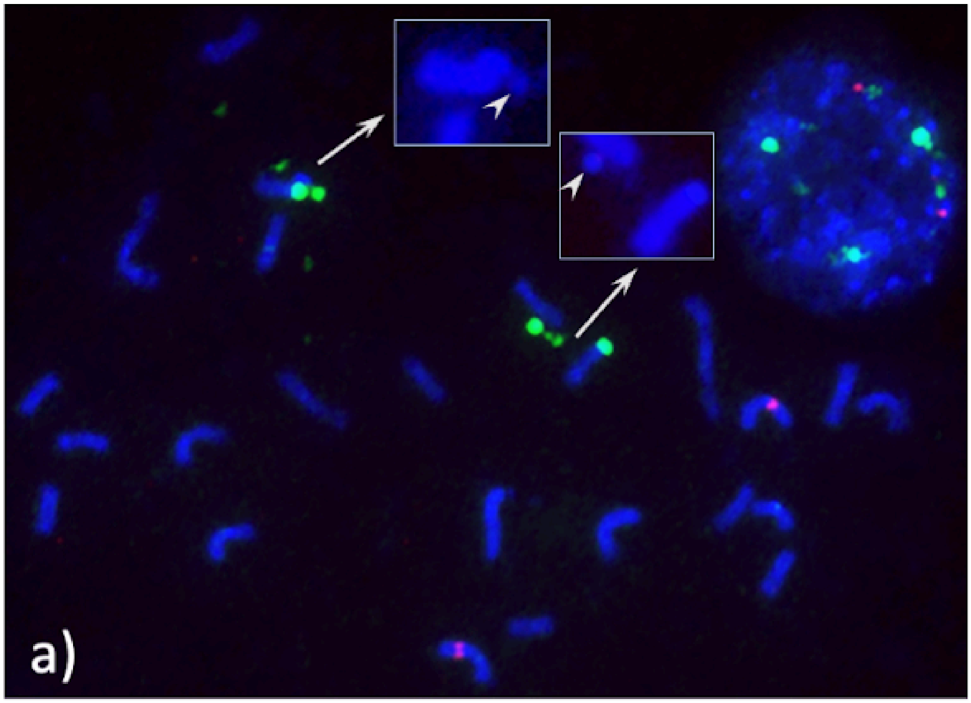

**Figure S6.** Major modes recovered with CLUMPAK for *Structure* results for K=1 to 5. Full output from CLUMPAK for all Structure analyses from K=1 to 10 is available from the data repository submission that accompanies this article.

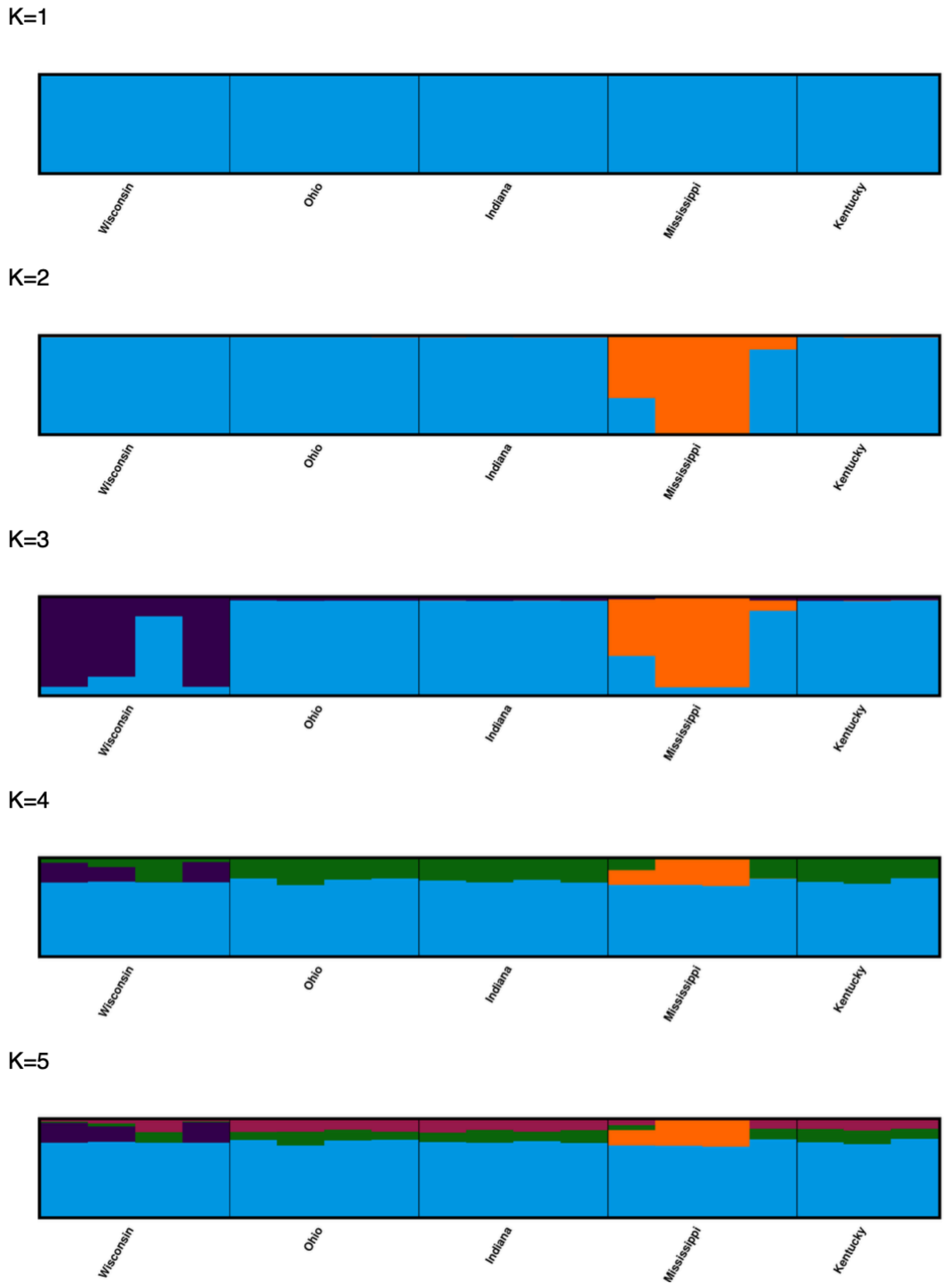

**Figure S7.** Phylogenetic tree of *Quercus* including all sampled individuals of *Quercus alba*. Colored boxes indicate clades (excluding *Q. alba*) used to calculate the number of variable sites shared with *Q. alba* in the phylogenomic matrix.

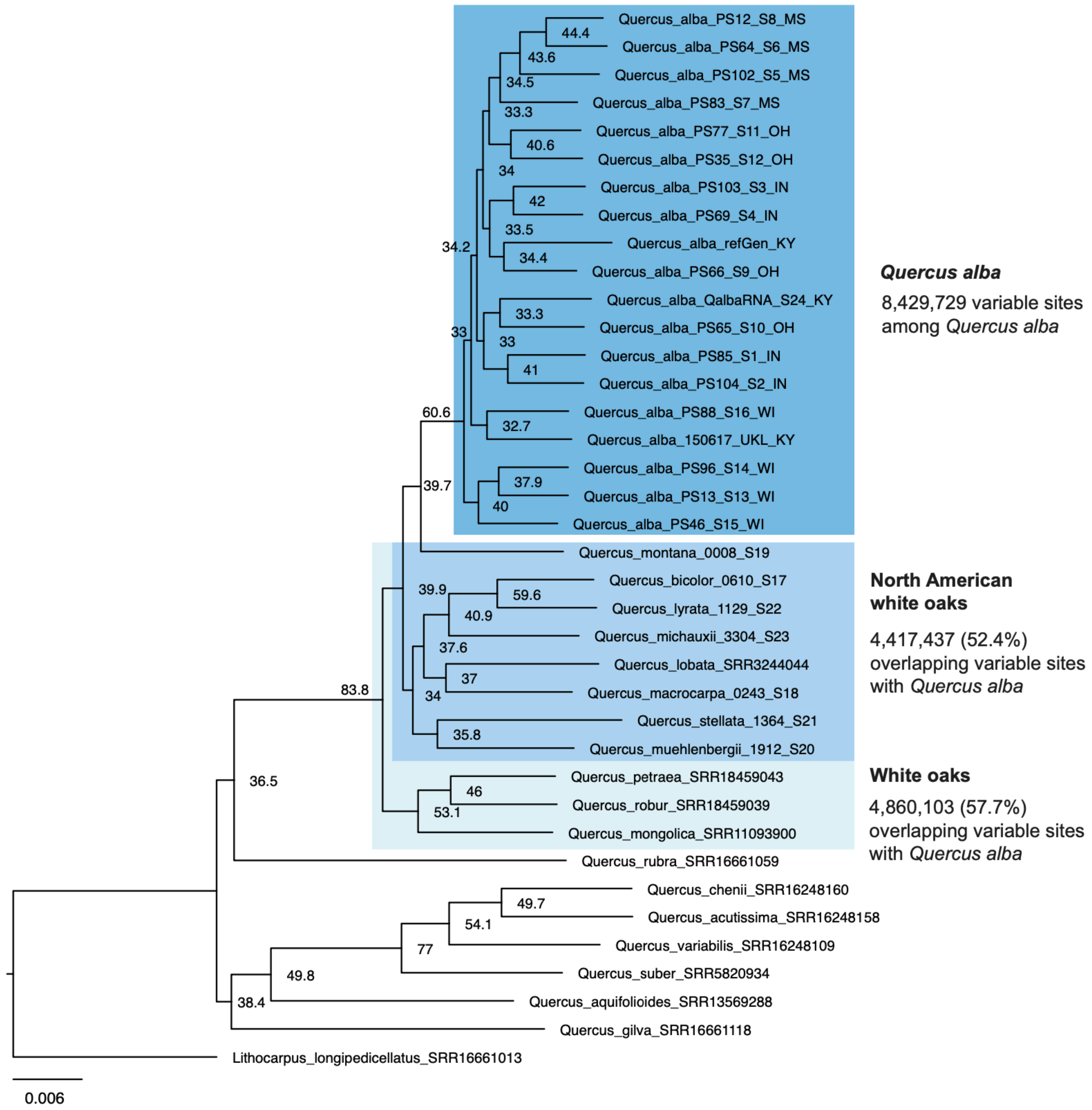

**Figure S8.** The ASTRAL tree, generated from 12,081 gene trees based on 5 kb windows. Node values are ASTRAL local posterior probabilities (left) and gene concordance factors (right). The tree is rooted on *Lithocarpus longipedicellatus*. Branch lengths are in coalescent units.

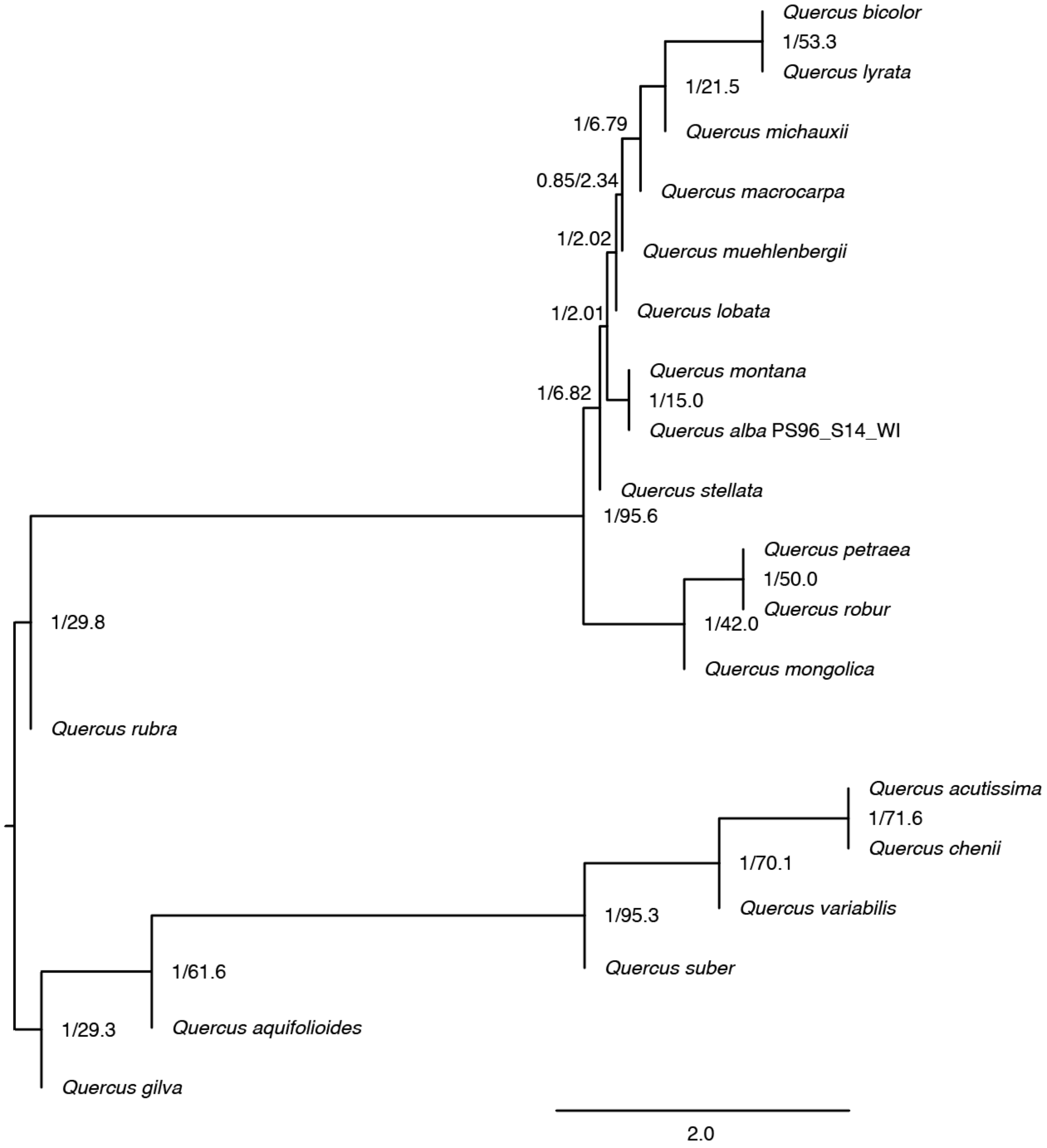

**Figure S9.** The plastome tree. Rooted on *Lithocarpus longipedicellatus*. Branch lengths are in units of inferred substitutions per site based on 93,946 aligned sites.

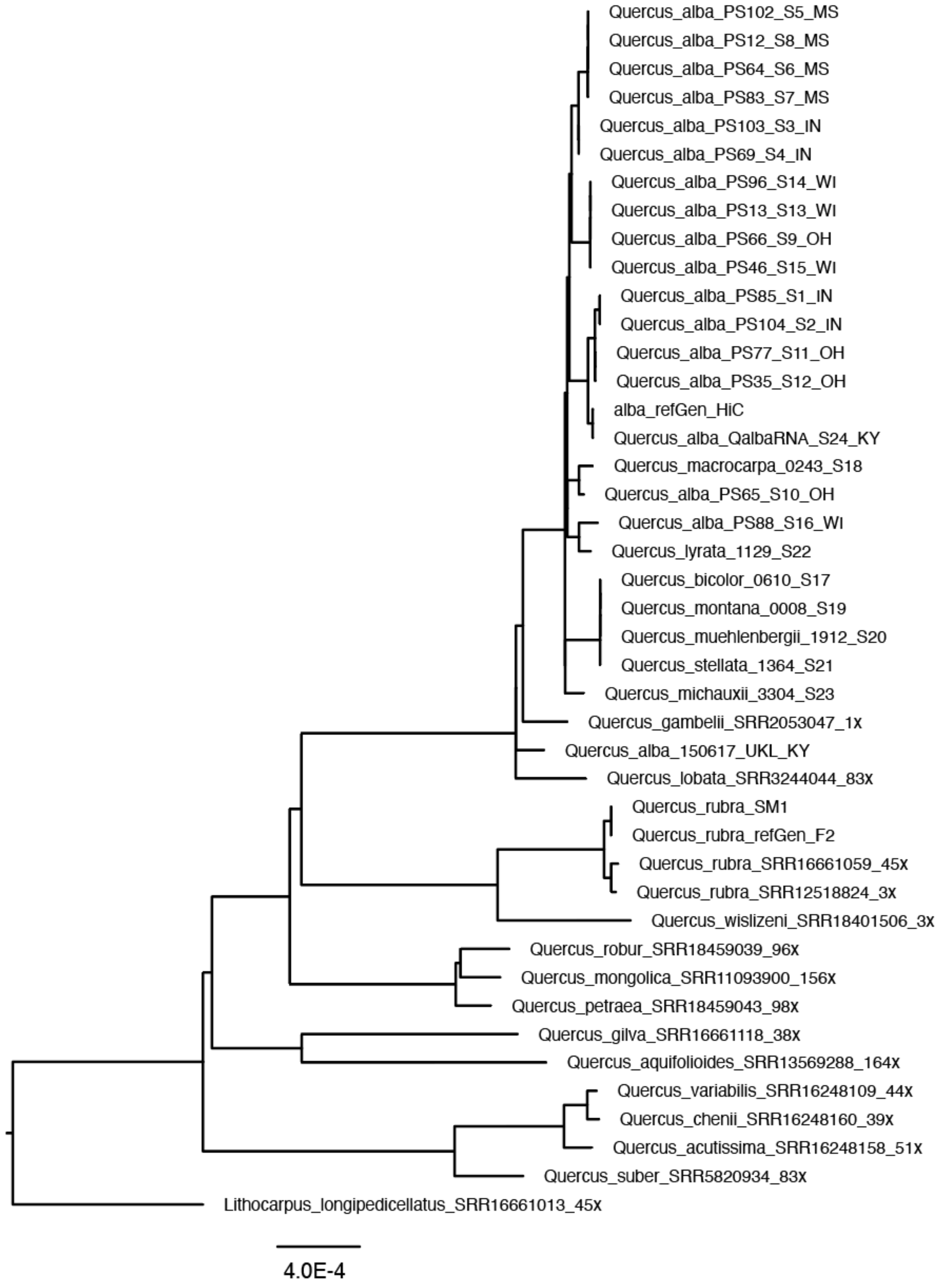

**Figure S10.** The mitochondrial genome tree. Rooted on *Lithocarpus longipedicellatus*. Branch lengths are in units of inferred substitutions per site based on 33,257 aligned sites.

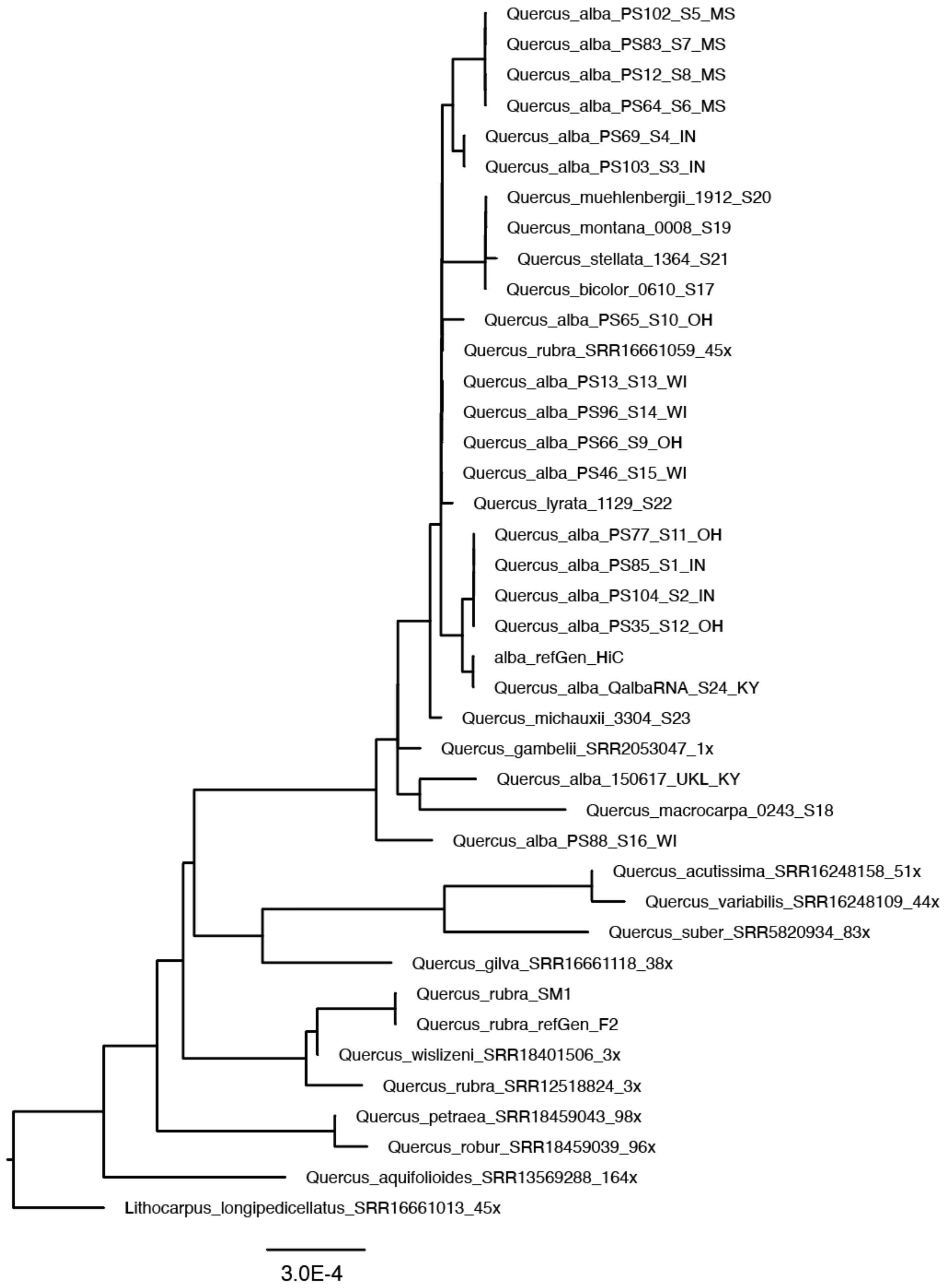

**Figure S11.** Topological comparison between the plastome (left) and mitogenome (right) trees. Both trees are rooted on *Lithocarpus* (not depicted).

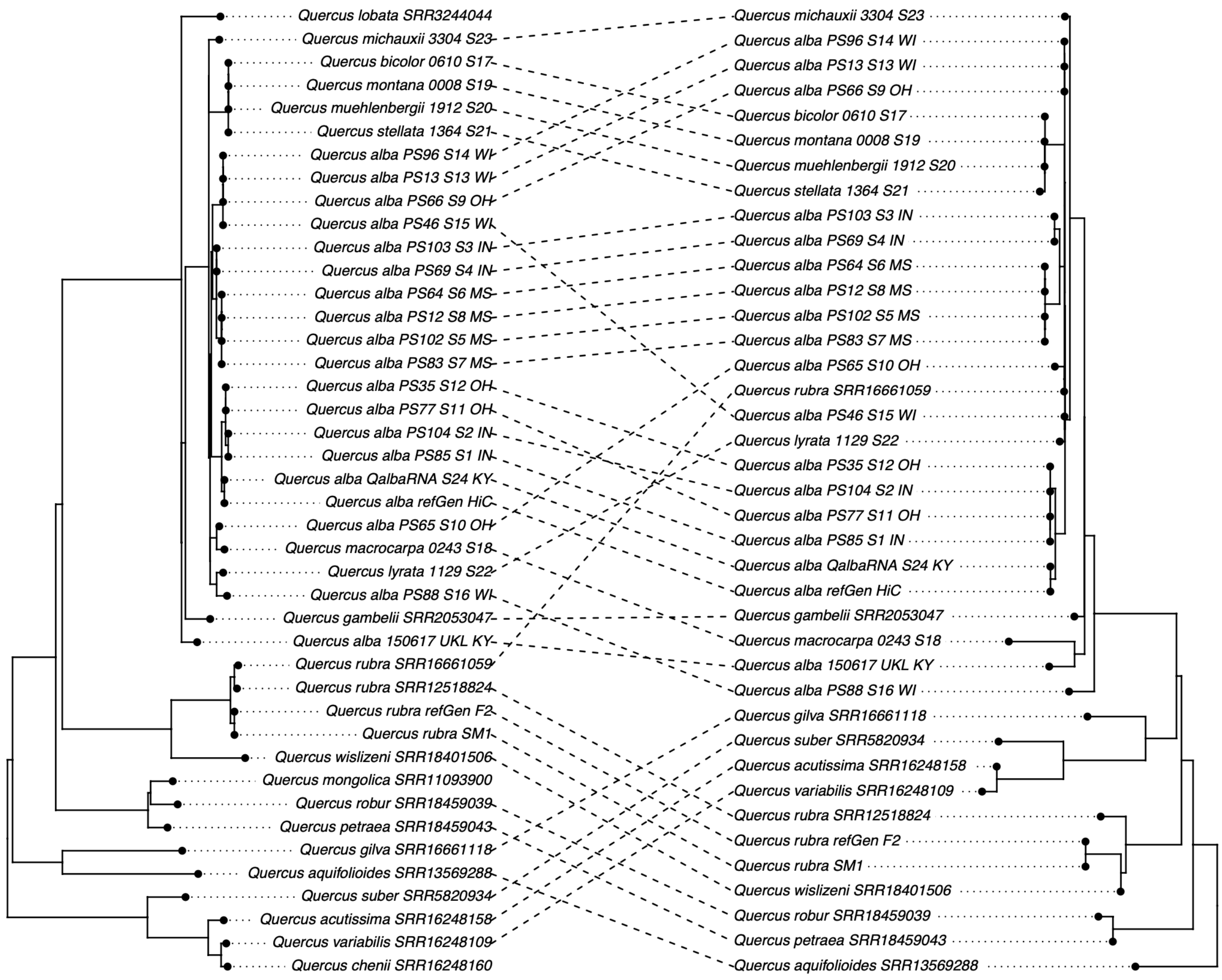

**Figure S12.** Dated phylogenies. Node labels are divergence times in millions of years since the present. A) Species divergence times after correction for within species nucleotide diversity. B) Allelic divergence times without correction for within species nucleotide diversity. *Crown age of *Quercus* was calibrated to 56 Ma.

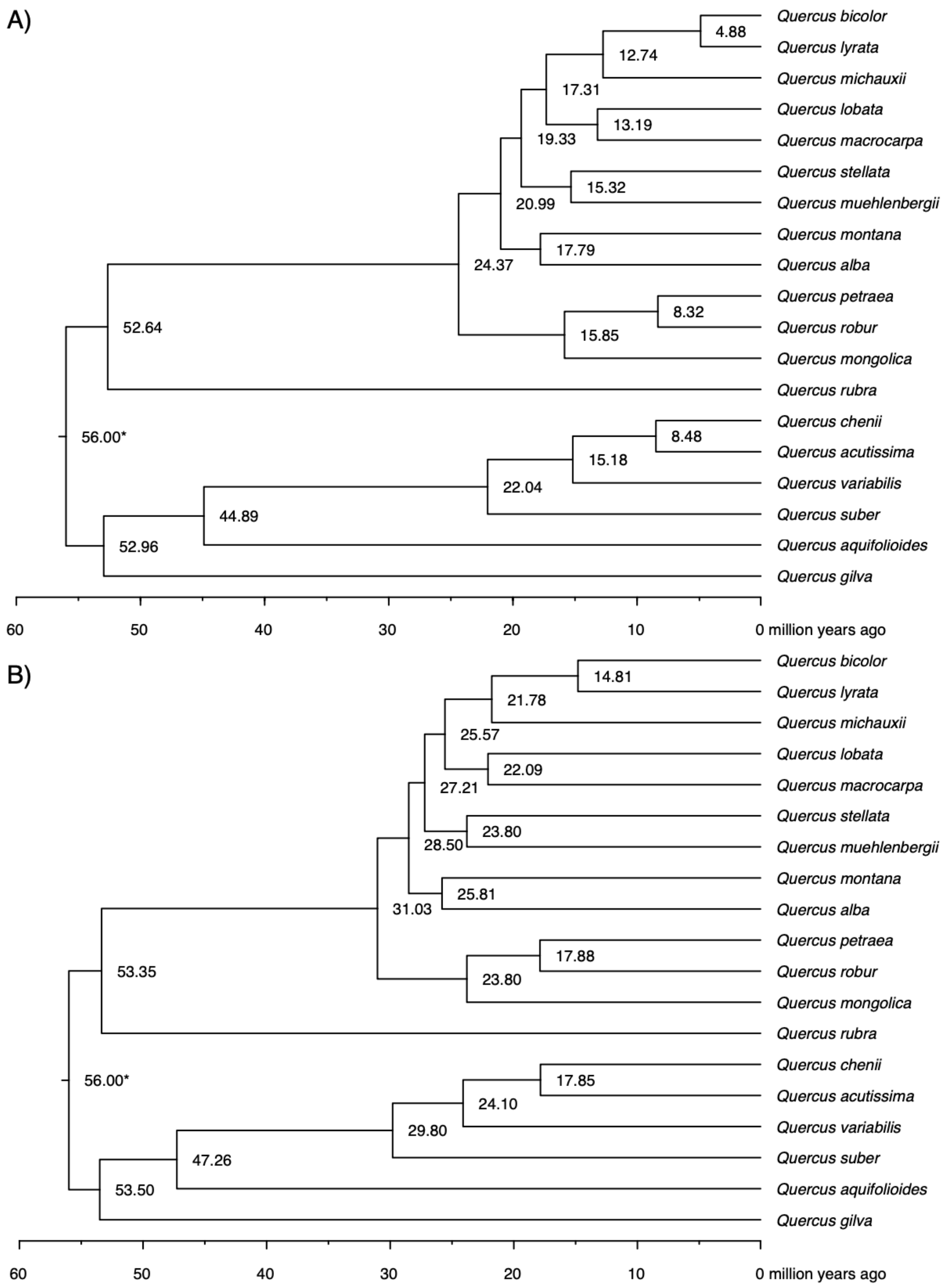

**Figure S13**. Overview of structural synteny of *Quercus* genomes, from top to bottom: *Q. rubra* (Qru), *Q. robur* (Qro), *Q. mongolica* (Qmo), *Q. lobata* (Qlo), *Q. gilva* (Qgi), *Q. alba* (Qal), *Q. acutissima* (Qac).

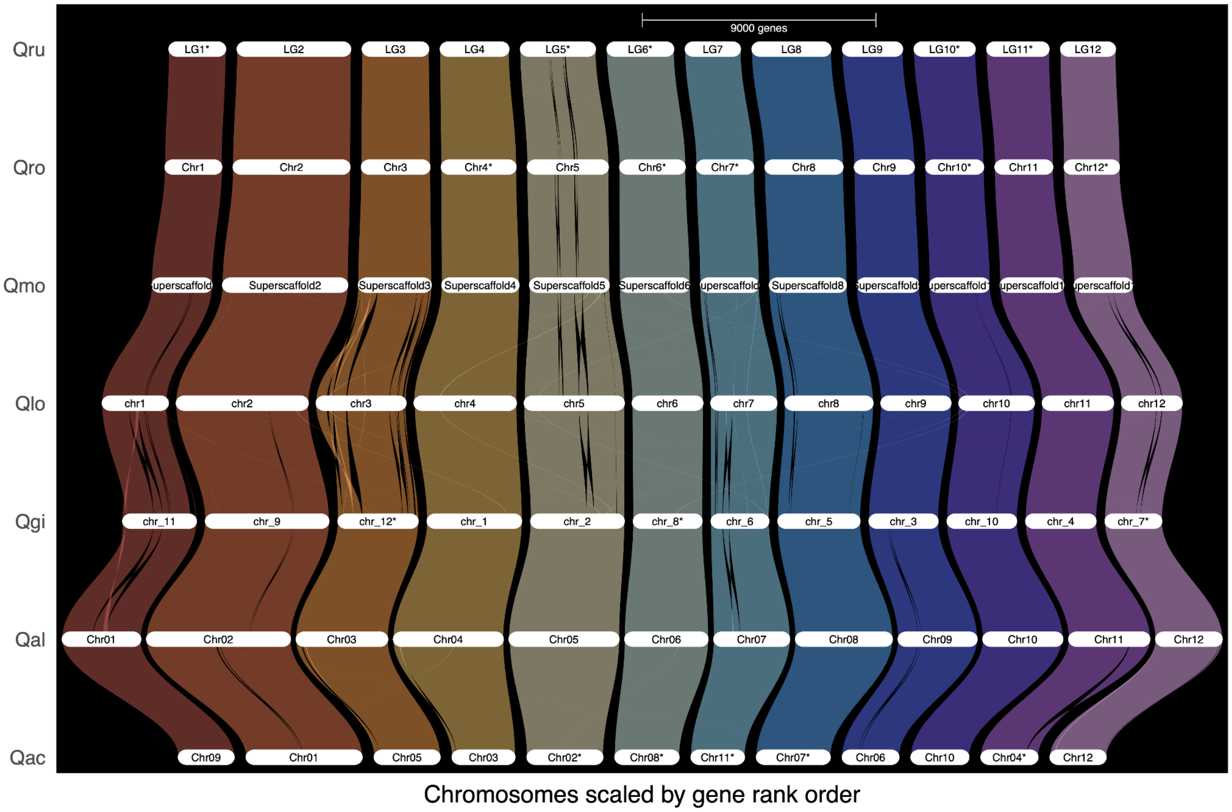

**Figure S14**. Detailed structural synteny of Q. alba Hap A against Q. mongolica (upper left), Q. lobata (upper right), Castanea mollissima (lower left), and Castanopsis tibetana (lower right).

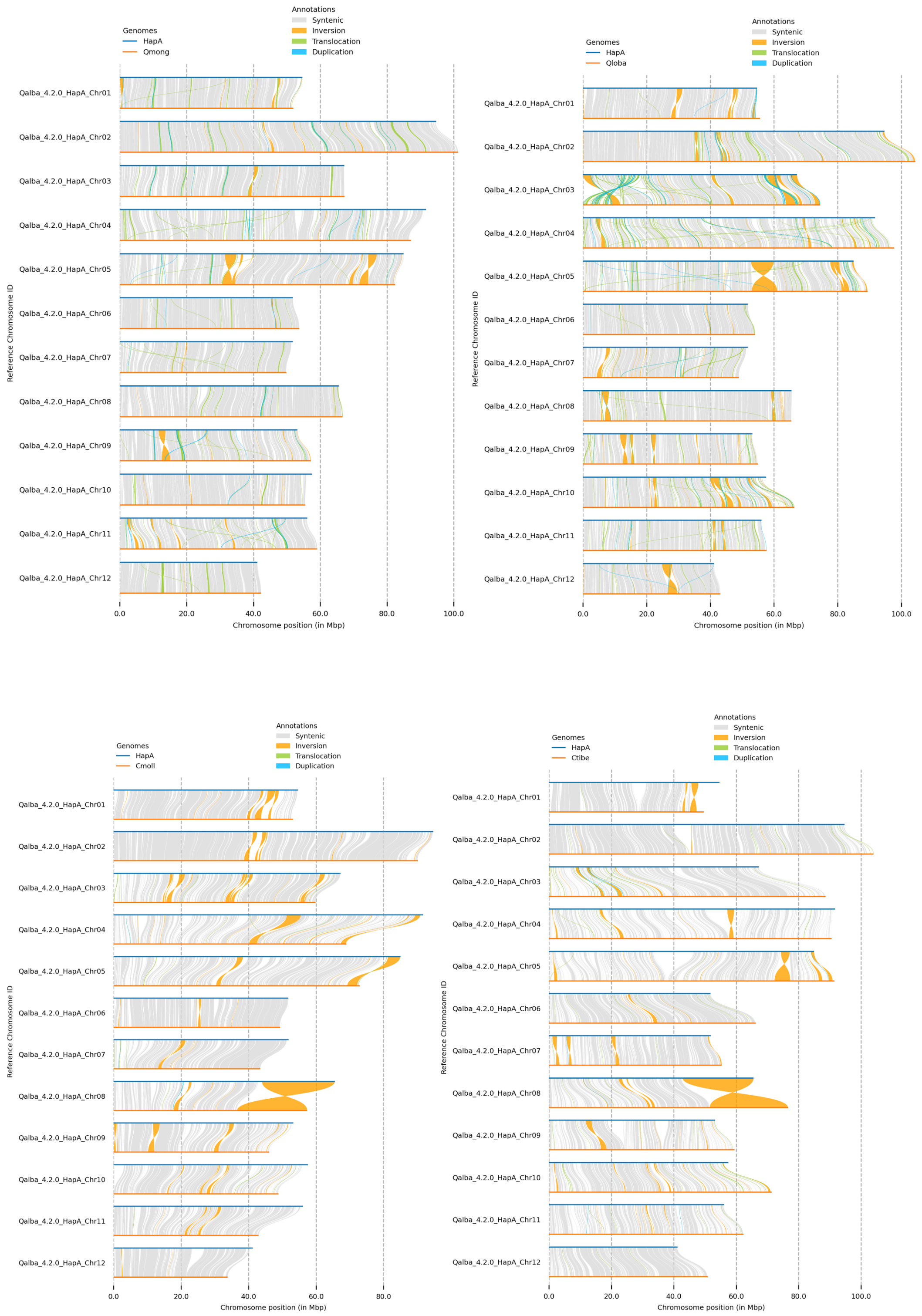

**Figure S15.** Gene duplicate classification of all genes and R genes from *Quercus* genomes.

A)

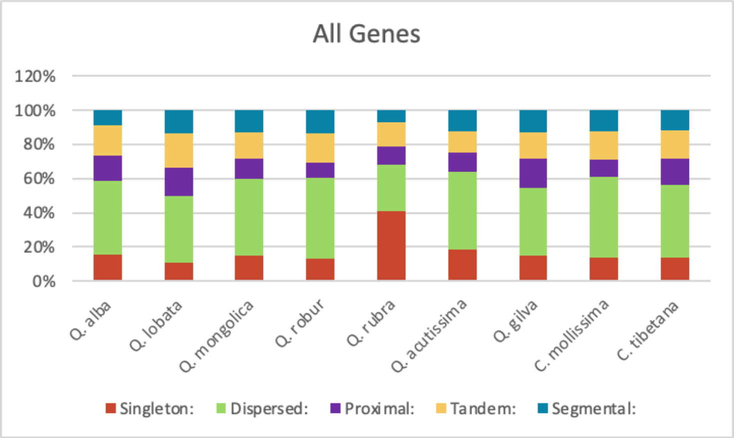


B)

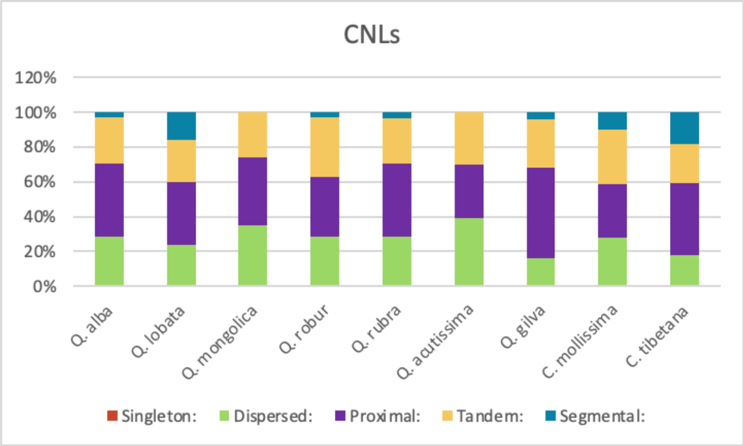


C)

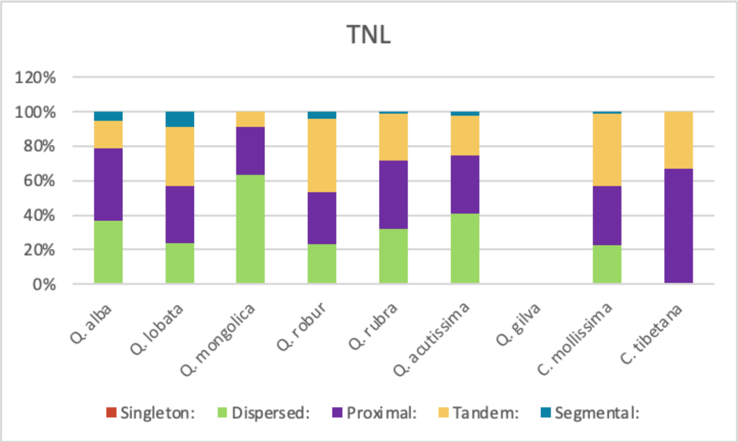


D)

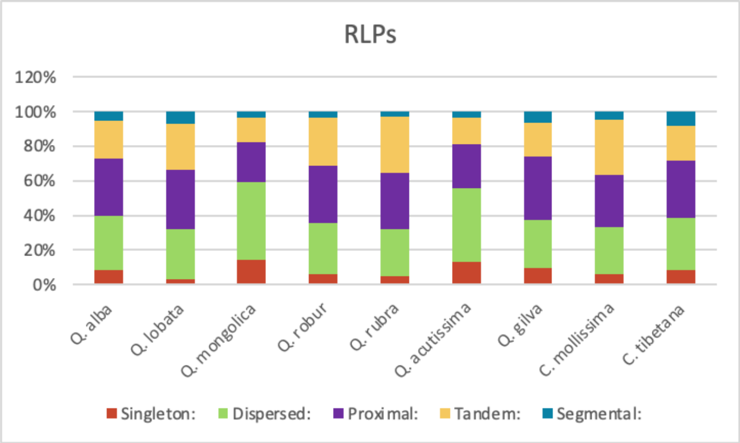


E)

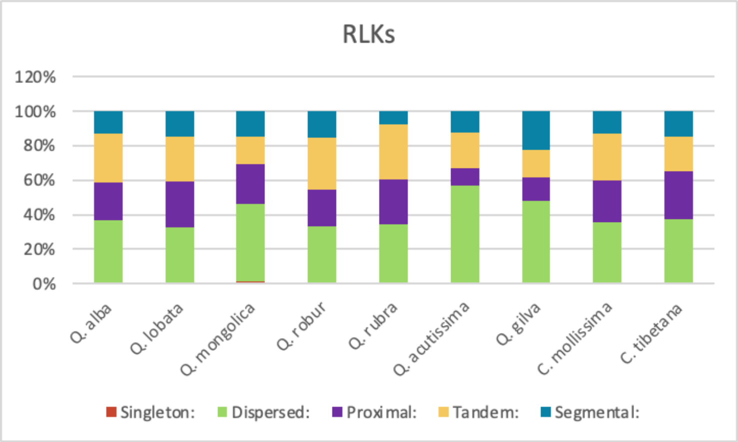


**Ai W, Liu Y, Mei M, Zhang X, Tan E, Liu H, Han X, Zhan H, Lu X**. **2022**. A chromosome‐scale genome assembly of the Mongolian oak ( *Quercus mongolica* ). *Molecular Ecology Resources* **22**: 2396–2410.

**Anders S, Pyl PT, Huber W**. **2015**. HTSeq—a Python framework to work with high-throughput sequencing data. *bioinformatics* **31**: 166–169.

**Andrews S**. **2010**. FastQC:  A Quality Control Tool for High Throughput Sequence Data.

**Bairoch A, Apweiler R, Wu CH, Barker WC, Boeckmann B, Ferro S, Gasteiger E, Huang H, Lopez R, Magrane M**. **2005**. The universal protein resource (UniProt). *Nucleic acids research* **33**: D154–D159.

**Broad Institute**. **2023**. Picard Tools.

**Brown JW, Walker JF, Smith SA**. **2017**. Phyx: phylogenetic tools for unix. *Bioinformatics (Oxford, England)* **33**: 1886–1888.

**Bruna T, Hoff KJ, Lomsadze A, Stanke M, Borodovsky M**. **2021**. BRAKER2: automatic eukaryotic genome annotation with GeneMark-EP+ and AUGUSTUS supported by a protein database. *NAR genomics and bioinformatics* **3**: lqaa108.

**Bruna T, Lomsadze A, Borodovsky M**. **2020**. GeneMark-EP+: eukaryotic gene prediction with self-training in the space of genes and proteins. *NAR genomics and bioinformatics* **2**: lqaa026.

**Caballero M, Wegrzyn J**. **2019**. gFACs: gene filtering, analysis, and conversion to unify genome annotations across alignment and gene prediction frameworks. *Genomics, proteomics & bioinformatics* **17**: 305–310.

**Cai L, Zhang H, Davis CC**. **2022**. PhyloHerb: A high‐throughput phylogenomic pipeline for processing genome skimming data. *Applications in Plant Sciences* **10**: e11475.
**Callahan AM, Zhebentyayeva TN, Humann JL, Saski CA, Galimba KD, Georgi LL, Scorza R, Main D, Dardick CD**. **2021**. Defining the 'HoneySweet'insertion event utilizing NextGen sequencing and a de novo genome assembly of plum (Prunus domestica). *Horticulture research* **8**.
**Calle García J, Guadagno A, Paytuvi-Gallart A, Saera-Vila A, Amoroso CG, D'Esposito D, Andolfo G, Aiese Cigliano R, Sanseverino W, Ercolano MR**. **2022**. PRGdb 4.0: an updated database dedicated to genes involved in plant disease resistance process. *Nucleic Acids Research* **50**: D1483–D1490.
**Camacho C, Coulouris G, Avagyan V, Ma N, Papadopoulos J, Bealer K, Madden TL**. **2009**. BLAST+: architecture and applications. *BMC Bioinformatics* **10**: 421.
**Cheng H, Concepcion GT, Feng X, Zhang H, Li H**. **2021**. Haplotype-resolved de novo assembly using phased assembly graphs with hifiasm. *Nature methods* **18**: 170–175.
**Danecek P, Auton A, Abecasis G, Albers CA, Banks E, DePristo MA, Handsaker RE, Lunter G, Marth GT, Sherry ST, *et al.*2011**. The variant call format and VCFtools. *Bioinformatics (Oxford, England)* **27**: 2156–2158.
**DePristo MA, Banks E, Poplin R, Garimella KV, Maguire JR, Hartl C, Philippakis AA, Del Angel G, Rivas MA, Hanna M**. **2011**. A framework for variation discovery and genotyping using next-generation DNA sequencing data. *Nature genetics* **43**: 491–498.
**Dobin A, Davis CA, Schlesinger F, Drenkow J, Zaleski C, Jha S, Batut P, Chaisson M, Gingeras TR**. **2013**. STAR: ultrafast universal RNA-seq aligner. *Bioinformatics* **29**: 15–21.
**Dudchenko O, Batra SS, Omer AD, Nyquist SK, Hoeger M, Durand NC, Shamim MS, Machol I, Lander ES, Aiden AP, *et al.*2017**. De novo assembly of the Aedes aegypti genome using Hi-C yields chromosome-length scaffolds. *Science (New York, N.Y.)* **356**: 92–95.
**Durand NC, Shamim MS, Machol I, Rao SSP, Huntley MH, Lander ES, Aiden EL**. **2016**. Juicer provides a one-click system for analyzing loop-resolution Hi-C experiments. *Cell systems* **3**: 95–98.
**Edwards SV, Beerli P**. **2000**. Perspective: gene divergence, population divergence, and the variance in coalescence time in phylogeographic studies. *Evolution; International Journal of Organic Evolution* **54**: 1839–1854.
**Ellinghaus D, Kurtz S, Willhoeft U**. **2008**. LTRharvest, an efficient and flexible software for de novo detection of LTR retrotransposons. *BMC Bioinformatics* **9**: 18.
**Emms DM, Kelly S**. **2019**. OrthoFinder: phylogenetic orthology inference for comparative genomics. *Genome Biology* **20**: 238.
**Feng Y**. **2003**. Plant MITEs: Useful Tools for Plant Genetics and Genomics. *Genomics, Proteomics & Bioinformatics* **1**: 90–100.
**Flynn JM, Hubley R, Goubert C, Rosen J, Clark AG, Feschotte C, Smit AF**. **2020**. RepeatModeler2 for automated genomic discovery of transposable element families. *Proceedings of the National Academy of Sciences* **117**: 9451–9457.
**Fu R, Zhu Y, Liu Y, Feng Y, Lu R-S, Li Y, Li P, Kremer A, Lascoux M, Chen J**. **2022**. Genome-wide analyses of introgression between two sympatric Asian oak species. *Nature Ecology & Evolution* **6**: 924–935.
**Gabriel L, Hoff KJ, Brůna T, Borodovsky M, Stanke M**. **2021**. TSEBRA: transcript selector for BRAKER. *BMC Bioinformatics* **22**: 566.
**Goel M, Schneeberger K**. **2022**. plotsr: visualizing structural similarities and rearrangements between multiple genomes. *Bioinformatics* **38**: 2922–2926.
**Goel M, Sun H, Jiao W-B, Schneeberger K**. **2019**. SyRI: finding genomic rearrangements and local sequence differences from whole-genome assemblies. *Genome Biology* **20**: 277.
**González J, Petrov D**. **2009**. MITEs—The Ultimate Parasites. *Science* **325**: 1352–1353.
**Greiner S, Lehwark P, Bock R**. **2019**. OrganellarGenomeDRAW (OGDRAW) version 1.3. 1: expanded toolkit for the graphical visualization of organellar genomes. *Nucleic acids research* **47**: W59–W64.
**Han B, Wang L, Xian Y, Xie X-M, Li W-Q, Zhao Y, Zhang R-G, Qin X, Li D-Z, Jia K-H**. **2022**. A chromosome-level genome assembly of the Chinese cork oak (Quercus variabilis). *Frontiers in Plant*

*Science* **13**: 1001583.
**Hao Z, Lv D, Ge Y, Shi J, Weijers D, Yu G, Chen J**. **2020**. RIdeogram: drawing SVG graphics to visualize and map genome-wide data on the idiograms. *PeerJ Computer Science* **6**: e251.
**Hart AJ, Ginzburg S, Xu M (Sam), Fisher CR, Rahmatpour N, Mitton JB, Paul R, Wegrzyn JL**. **2020**. EnTAP: Bringing faster and smarter functional annotation to non‐model eukaryotic transcriptomes. *Molecular Ecology Resources* **20**: 591–604.
**Hipp AL, Manos PS, Hahn M, Avishai M, Bodénès C, Cavender‐Bares J, Crowl AA, Deng M, Denk T, Fitz‐Gibbon S, *et al.*2020**. Genomic landscape of the global oak phylogeny. *New Phytologist* **226**: 1198–1212.
**Hofmann C-C**. **2010**. Microstructure of Fagaceae pollen from Austria (Palaeocene/Eocene boundary) and Hainan Island, (?Middle Eocene) 8 th European Palaeobotany and Palynology Conference 2010 in Budapest.
**Hofmann C-Ch, Mohamed O, Egger H**. **2011**. A new terrestrial palynoflora from the Palaeocene/Eocene boundary in the northwestern Tethyan realm (St. Pankraz, Austria). *Review of Palaeobotany and Palynology* **166**: 295–310.
**Hu H-L, Zhang J-Y, Li Y-P, Xie L, Chen D-B, Li Q, Liu Y-Q, Hui S-R, Qin L**. **2019**. The complete chloroplast genome of the daimyo oak, Quercus dentata Thunb. *Conservation Genetics Resources* **11**: 409–411.
**Hubley R, Finn RD, Clements J, Eddy SR, Jones TA, Bao W, Smit AF, Wheeler TJ**. **2016**. The Dfam database of repetitive DNA families. *Nucleic acids research* **44**: D81–D89.
**Hudson RR, Slatkin M, Maddison WP**. **1992**. Estimation of levels of gene flow from DNA sequence data. *Genetics* **132**: 583–589.
**International Peach Genome Initiative, Verde I, Abbott AG, Scalabrin S, Jung S, Shu S, Marroni F, Zhebentyayeva T, Dettori MT, Grimwood J, *et al.*2013**. The high-quality draft genome of peach (Prunus persica) identifies unique patterns of genetic diversity, domestication and genome evolution. *Nature Genetics* **45**: 487–494.
**Jia H-M, Jia H-J, Cai Q-L, Wang Y, Zhao H-B, Yang W-F, Wang G-Y, Li Y-H, Zhan D-L, Shen Y-T, *et al.*2019**. The red bayberry genome and genetic basis of sex determination. *Plant Biotechnology Journal* **17**: 397–409.
**Jiang H, Lei R, Ding S-W, Zhu S**. **2014**. Skewer: a fast and accurate adapter trimmer for next-generation sequencing paired-end reads. *BMC Bioinformatics* **15**: 182.
**Kapoor B, Jenkins J, Schmutz J, Zhebentyayeva T, Kuelheim C, Coggeshall M, Heim C, Lasky JR, Leites L, Islam-Faridi N**. **2023**. A haplotype-resolved chromosome-scale genome for Quercus rubra L. provides insights into the genetics of adaptive traits for red oak species. *G3: Genes, Genomes, Genetics* **13**: jkad209.
**Katoh K, Misawa K, Kuma K, Miyata T**. **2002**. MAFFT: a novel method for rapid multiple sequence alignment based on fast Fourier transform. *Nucleic acids research* **30**: 3059–3066.
**Katoh K, Standley DM**. **2013**. MAFFT multiple sequence alignment software version 7: improvements in performance and usability. *Molecular biology and evolution* **30**: 772–780.
**Kopelman NM, Mayzel J, Jakobsson M, Rosenberg NA, Mayrose I**. **2015**. Clumpak: a program for identifying clustering modes and packaging population structure inferences across K. *Molecular Ecology Resources* **15**: 1179–1191.
**Lagesen K, Hallin P, Rødland EA, Stærfeldt H-H, Rognes T, Ussery DW**. **2007**. RNAmmer: consistent and rapid annotation of ribosomal RNA genes. *Nucleic acids research* **35**: 3100–3108.
**Larsson A**. **2014**. AliView: a fast and lightweight alignment viewer and editor for large datasets. *Bioinformatics* **30**: 3276–3278.
**Li H**. **2011**. A statistical framework for SNP calling, mutation discovery, association mapping and population genetical parameter estimation from sequencing data. *Bioinformatics (Oxford, England)* **27**: 2987–2993.
**Li H**. **2018**. Minimap2: pairwise alignment for nucleotide sequences. *Bioinformatics* **34**: 3094–3100.
**Li H, Durbin R**. **2009**. Fast and accurate short read alignment with Burrows-Wheeler transform.

*Bioinformatics* **25**: 1754–1760.
**Li H, Handsaker B, Wysoker A, Fennell T, Ruan J, Homer N, Marth G, Abecasis G, Durbin R, 1000 Genome Project Data Processing Subgroup**. **2009**. The Sequence Alignment/Map format and SAMtools. *Bioinformatics* **25**: 2078–2079.
**Lovell JT, Bentley NB, Bhattarai G, Jenkins JW, Sreedasyam A, Alarcon Y, Bock C, Boston LB, Carlson J, Cervantes K, *et al.*2021**. Four chromosome scale genomes and a pan-genome annotation to accelerate pecan tree breeding. *Nature Communications* **12**: 4125.
**Lovell JT, Sreedasyam A, Schranz ME, Wilson M, Carlson JW, Harkess A, Emms D, Goodstein DM, Schmutz J**. **2022**. GENESPACE tracks regions of interest and gene copy number variation across multiple genomes. *eLife* **11**: e78526.
**Lucas SJ, Kahraman K, Avşar B, Buggs RJA, Bilge I**. **2021**. A chromosome-scale genome assembly of European hazel (Corylus avellana L.) reveals targets for crop improvement. *The Plant Journal: For Cell and Molecular Biology* **105**: 1413–1430.
**Manni M, Berkeley MR, Seppey M, Simão FA, Zdobnov EM**. **2021**. BUSCO Update: Novel and Streamlined Workflows along with Broader and Deeper Phylogenetic Coverage for Scoring of Eukaryotic, Prokaryotic, and Viral Genomes. *Molecular Biology and Evolution* **38**: 4647–4654.
**Martínez-García PJ, Crepeau MW, Puiu D, Gonzalez-Ibeas D, Whalen J, Stevens KA, Paul R, Butterfield TS, Britton MT, Reagan RL, *et al.*2016**. The walnut (Juglans regia) genome sequence reveals diversity in genes coding for the biosynthesis of non-structural polyphenols. *The Plant Journal: For Cell and Molecular Biology* **87**: 507–532.
**McKenna A, Hanna M, Banks E, Sivachenko A, Cibulskis K, Kernytsky A, Garimella K, Altshuler D, Gabriel S, Daly M**. **2010**. The Genome Analysis Toolkit: a MapReduce framework for analyzing next-generation DNA sequencing data. *Genome research* **20**: 1297–1303.
**Mendes FK, Vanderpool D, Fulton B, Hahn MW**. **2020**. CAFE 5 models variation in evolutionary rates among gene families. *Bioinformatics (Oxford, England)* **36**: 5516–5518.
**Minh BQ, Hahn MW, Lanfear R**. **2020a**. New Methods to Calculate Concordance Factors for Phylogenomic Datasets. *Molecular Biology and Evolution* **37**: 2727–2733.
**Minh BQ, Schmidt HA, Chernomor O, Schrempf D, Woodhams MD, von Haeseler A, Lanfear R**. **2020b**. IQ-TREE 2: New Models and Efficient Methods for Phylogenetic Inference in the Genomic Era. *Molecular Biology and Evolution* **37**: 1530–1534.
**Mishra B, Ulaszewski B, Meger J, Aury J-M, Bodénès C, Lesur-Kupin I, Pfenninger M, Da Silva C, Gupta DK, Guichoux E, *et al.*2021**. A Chromosome-Level Genome Assembly of the European Beech (Fagus sylvatica) Reveals Anomalies for Organelle DNA Integration, Repeat Content and Distribution of SNPs. *Frontiers in Genetics* **12**: 691058.
**NCBI**. **2023**. SRA-tools.
**Ou S, Jiang N**. **2018**. LTR_retriever: a highly accurate and sensitive program for identification of long terminal repeat retrotransposons. *Plant physiology* **176**: 1410–1422.
**Paradis E, Schliep K**. **2019**. ape 5.0: an environment for modern phylogenetics and evolutionary analyses in R. *Bioinformatics* **35**: 526–528.
**Pritchard JK, Stephens M, Donnelly P**. **2000**. Inference of population structure using multilocus genotype data. *Genetics* **155**: 945–959.
**Purcell S, Neale B, Todd-Brown K, Thomas L, Ferreira MA, Bender D, Maller J, Sklar P, De Bakker PI, Daly MJ**. **2007**. PLINK: a tool set for whole-genome association and population-based linkage analyses. *The American journal of human genetics* **81**: 559–575.
**Quinlan AR, Hall IM**. **2010**. BEDTools: a flexible suite of utilities for comparing genomic features. *Bioinformatics* **26**: 841–842.
**R Core Team R**. **2013**. R: A language and environment for statistical computing.
**Revell LJ**. **2012**. phytools: an R package for phylogenetic comparative biology (and other things). *Methods in Ecology and Evolution* **3**: 217–223.
**Robinson DF, Foulds LR**. **1981**. Comparison of phylogenetic trees. *Mathematical biosciences* **53**: 131–147.

**Robinson JT, Turner D, Durand NC, Thorvaldsdóttir H, Mesirov JP, Aiden EL**. **2018**. Juicebox.js Provides a Cloud-Based Visualization System for Hi-C Data. *Cell systems* **6**: 256-258.e1.
**Sanderson MJ**. **2002**. Estimating absolute rates of molecular evolution and divergence times: a penalized likelihood approach. *Molecular Biology and Evolution* **19**: 101–109.
**Sim SB, Corpuz RL, Simmonds TJ, Geib SM**. **2022**. HiFiAdapterFilt, a memory efficient read processing pipeline, prevents occurrence of adapter sequence in PacBio HiFi reads and their negative impacts on genome assembly. *BMC Genomics* **23**: 157.
**Smit A, Hubley R, Green P**. **2004**. RepeatMasker Open-4.0.
**Smith SA, O'Meara BC**. **2012**. treePL: divergence time estimation using penalized likelihood for large phylogenies. *Bioinformatics (Oxford, England)* **28**: 2689–2690.
**Soltani N, Best T, Grace D, Nelms C, Shumaker K, Romero-Severson J, Moses D, Schuster S, Staton M, Carlson J, *et al.*2020**. Transcriptome profiles of Quercus rubra responding to increased O3 stress. *BMC Genomics* **21**: 160.
**Sork VL, Cokus SJ, Fitz-Gibbon ST, Zimin AV, Puiu D, Garcia JA, Gugger PF, Henriquez CL, Zhen Y, Lohmueller KE**. **2022**. High-quality genome and methylomes illustrate features underlying evolutionary success of oaks. *Nature communications* **13**: 2047.
**Sun Y, Guo J, Zeng X, Chen R, Feng Y, Chen S, Yang K**. **2022**. Chromosome-scale genome assembly of Castanopsis tibetana provides a powerful comparative framework to study the evolution and adaptation of Fagaceae trees. *Molecular Ecology Resources* **22**: 1178–1189.
**The Darwin Tree of Life Project Consortium, Blaxter M, Mieszkowska N, Di Palma F, Holland P, Durbin R, Richards T, Berriman M, Kersey P, Hollingsworth P, *et al.*2022**. Sequence locally, think globally: The Darwin Tree of Life Project. *Proceedings of the National Academy of Sciences* **119**: e2115642118.
**Thomas GWC, Hahn MW**. **2019**. Referee: Reference Assembly Quality Scores. *Genome Biology and Evolution* **11**: 1483–1486.
**Van Der Auwera GA, Carneiro MO, Hartl C, Poplin R, Del Angel G, Levy‐Moonshine A, Jordan T, Shakir K, Roazen D, Thibault J, *et al.***2013. From FastQ Data to High‐Confidence Variant Calls: The Genome Analysis Toolkit Best Practices Pipeline. *Current Protocols in Bioinformatics* **43**.
**Wang Y, Tang H, DeBarry JD, Tan X, Li J, Wang X, Lee T, Jin H, Marler B, Guo H**. **2012**. MCScanX: a toolkit for detection and evolutionary analysis of gene synteny and collinearity. *Nucleic acids research* **40**: e49–e49.
**Wang J, Tian S, Sun X, Cheng X, Duan N, Tao J, Shen G**. **2020**. Construction of Pseudomolecules for the Chinese Chestnut (Castanea mollissima) Genome. *G3 (Bethesda, Md.)* **10**: 3565–3574.
**Weir BS, Cockerham CC**. **1984**. ESTIMATING F-STATISTICS FOR THE ANALYSIS OF POPULATION STRUCTURE. *Evolution; International Journal of Organic Evolution* **38**: 1358–1370.
**Wickham H**. **2016**. *ggplot2*. Springer-Verlag New York.
**Xu Z, Wang H**. **2007**. LTR_FINDER: an efficient tool for the prediction of full-length LTR retrotransposons. *Nucleic acids research* **35**: W265–W268.
**Yang X, Wang Z, Zhang L, Hao G, Liu J, Yang Y**. **2020**. A chromosome-level reference genome of the hornbeam, Carpinus fangiana. *Scientific Data* **7**: 24.
**Zhou X, Liu N, Jiang X, Qin Z, Farooq TH, Cao F, Li H**. **2022**. A chromosome-scale genome assembly of Quercus gilva: Insights into the evolution of Quercus section Cyclobalanopsis (Fagaceae). *Frontiers in Plant Science* **13**: 1012277.
